# dCas9 targeted proteome profiling reveals p300-mediated reciprocal regulation of SMAD and SP1 as a driver of GM2synthase transcription in renal cell carcinoma

**DOI:** 10.1101/2025.09.16.676212

**Authors:** Sounak Banerjee, Avisek Banerjee, Subha Ray, Aishwarya Ray, Debarati Paul, Shubhra Ghosh Dastidar, Belinda Willard, Kaushik Biswas

**Affiliations:** Department of Biological Sciences, Bose Institute, Kolkata, West Bengal, 700091, India; Department of Zoology, Ramakrishna Mission Vidyamandira, Belur Math, Howrah, 711202, India; Proteomics and Metabolomics SLR, Lerner Research Institute, Cleveland Clinic, Cleveland, OH, USA, 44195

**Author notes:** Kaushik Biswas, Department of Biological Sciences, Bose Institute, Kolkata, EN-80, Bidhan Nagar, Salt Lake, Sector-V, Kolkata, West Bengal, 700091, India. To whom all correspondence should be addressed (Kaushik Biswas). These authors have contributed equally to this work (Avisek Banerjee, Subha Ray).

**Keywords:** **Keyword:** dCas9-CLASP-WB, enChIP-MS, Epigenetic gene regulation, SP1, HAT-p300, SMAD

## Abstract

Glycolipids constitute an important component of the plasma membrane based on both abundance as well as function. Gangliosides, being a class of structurally diverse and functionally varied glycolipids, can act both as a receptor as well as a ligand and therefore is established as a crucial player in several normal cellular processes. In certain diseases, and in particular cancer, select gangliosides are over-expressed often leading to disease manifestation. *GM2-synthase*, the enzyme responsible for the formation of a pro-tumorigenic ganglioside, GM2 is well reported to be over-expressed across various cancer tissues and cell lines. This over-expression of *GM2-synthase* has been linked with increased migration, invasion and epithelial to mesenchymal transition (1) as well as induction of a local and systemic host immune suppression in cancer. Despite only a handful of studies demonstrating an epigenetic regulation underlying the transcriptional regulation of *GM2-synthase (B4GalNT1)* gene, the detailed mechanism still remains unclear. Here we identified the total proteome associated with the *GM2-synthase* promoter through a two-step CRISPR-dCas9 based proteome profiling approach by categorizing all the identified proteins leading to a detailed elucidation of the molecular drivers behind *GM2-synthase* transcription. While the previous study identified an acetylation-dependent de-repression of the transcription factor SP1 causing *GM2-synthase* activation, the underlying molecular mechanism driving its activation wasn’t clear. This study demonstrated that the histone acetyl transferase (2), p300 acts as a pivotal factor which on one hand cause acetylation-mediated degradation of SP1, and on the other hand activates SMAD2/4 to have a direct positive impact on *GM2-synthase* gene transcription. We identified p300 to have an activator role in *GM2-synthase* gene transcription through knock out, knock down and over-expression experiments. Furthermore, SP1 degradation, SMAD activation and their DNA binding patterns show the reciprocal role of p300 on SP1 and SMAD complexes. Altogether we have identified SMAD 2/4 as an activator complex, p300 as a positive regulator and uncovered a critical p300-SMAD-SP1 regulatory axis in *GM2-synthase* transcriptional regulation.

## Introduction

Cell surface lipids hold tremendous importance in biological systems as they form an integral part of the plasma membrane and perform critical cellular functions. Gangliosides are one such sialic acid containing glycosphingolipids residing on the outer layer of plasma membrane and are involved in normal cellular functions not limiting to formation of lipid rafts (3), cell-cell communication and cellular signaling (4). Aberrant over-expression of gangliosides and their corresponding glycosyltransferases were reported in various cancers (5–7). These cancer-specific over-expression have often been found to be associated with modulation of the tumor microenvironment including local as well as systemic immune suppression (5, 8). Several studies from the past have also established the role of gangliosides in modulating various aspects of tumor cell behavior (9–11). Research from our laboratory established ganglioside GM2 as one of the predominant tumor over-expressed gangliosides involved in promoting migration and invasion (2), imparting anoikis resistance thereby causing anchorage independence (AIG), overall shifting the cellular landscape from an epithelial to a mesenchymal phenotype characteristic of EMT (12) in cancer cells. Following this, *Sasaki et.al* (13) also showed ganglioside GM2 to be involved in regulation of growth and invasion of pancreatic ductal adenocarcinoma (PDAC). Despite emerging evidence indicating *GM2/GD2-synthase* over-expression to be involved in pathogenesis of select cancers, the precise molecular mechanisms underlying its over-expression remains enigmatic. Data from the Human Protein Atlas database shows *GM2-synthase* (GalNAcT, β4GalNT1) mRNA levels to be elevated in several tumor tissues (5–7), while that from the TCGA database linked this over-expression of *GM2-synthase* with poor prognosis in terms of overall survival (14), which prompted us to explore and identify the mechanisms behind the regulation of *GM2-synthase* transcription which still remains elusive in cancer.

Initial investigations about ganglioside synthase gene promoters revealed them as TATA-less, lacking a CCAAT box with a predominance of GC rich regions and having binding sites for transcription factors such as SP1 or AP-2 in the proximal promoter regions as some of the pertinent characteristics. However, their functional significance remained unclear (15). The genetic characteristics of β4GalNT1 or *GM2/GD2-synthase* gene and its regulatory elements were first uncovered by *Furukawa et.al* which revealed that the gene has 11 exons with three transcript variants spanning over a genomic length of over 8kb. It also assigned the genomic location of the gene to 12q13.3 along with a detailed profiling of the 5’- flanking region of exon1b with identification of several transcription factor binding elements present (16). The first evidence of an epigenetic mechanism regulating the transcription of the *GM2-synthase* was shown by *Suzuki et.al* (17), where histone acetylation was identified as a key mechanism regulating the expression of *GM2-synthase* gene at various mouse brain development stages. Immediately following this, our laboratory showed that *GM2-synthase* gene was not only epigenetically regulated by histone acetylation marks such as H3K9 and H3K14 acetylation at the gene’s TSS, but also how these marks are regulated by an intricate interplay between several DNA binding (such as SP1) and non-DNA binding (such as HDAC1) regulatory elements. We also established SP1-HDAC1 complex to act as a co-repressor complex for the transcriptional regulation of the *GM2-synthase* gene and established an acetylation mark at the K703 position of SP1 to be the critical factor between transcriptional repression and de-repression balance of *GM2-synthase* gene in cancer (18).

In the present study we not only mapped the molecular machinery involved but also identified a key regulatory switch that dictates the outcome of *GM2-synthase* transcription in high versus low *GM2-synthase* transcriptional states. The involvement of plausible non-DNA binding elements the binding of which to the DNA unlike direct DNA binding elements, cannot be efficiently predicted using bioinformatic tools, prompted us to profile the proteome associated with the *GM2-synthase* promoter and thereby identify critical players involved in the transcriptional process. Using the ability of CRISPR-Cas system to specifically target a desired locus on the genome, coupled with the *de-novo* identification power of mass spectrometry and the highly specific detection power of classical analytical methods like western blotting, we were able to successfully identify the proteome associated with the *GM2-synthase* promoter. We established SMAD 2/4 complex as an activator of transcription of the *GM2-synthase* gene. One of the unique findings of this study is the identification of a key switch, p300 (*EP300*) which by its ability to acetylate the transcriptional repressor, SP1 or modify and control the activator, SMAD2/4 dictates *GM2-synthase* transcriptional outcome.

## Materials and Methods

### Cell culture and transfection

NCCS, Pune, India provided HEK-293T (RRID:CVCL_0063) cells, while SK-RC-45 cells (RRID:CVCL_4016) were gifted by Dr. James H. Finke (Cleveland Clinic). HEK-293T cell line was cultured with DMEM (Gibco, CAT: 12800017) while SK-RC-45 cells were cultured using RPMI-1640 (Gibco, CAT:31800022) using standard procedures (2). Indicated plasmids were transfected to the cells using Lipofectamine-LTX (Life Technologies, Cat:15338100). Human recombinant TGFβ protein was procured from R&D Biosystems (Cat: 7754-BH-005/CF). For the experiments involving drug treatments, the drugs that were used are the following: Sodium butyrate (Sisco Research Laboratories, India, Cat: 22308), epigallocatechin-3-gallate (Sigma-Aldrich, Cat: 989-51-5). The HEK-293T cell line was obtained from NCCS, Pune, India and is source authenticated by STR profiling, while the SK-RC-45 cell line was received as a kind gift from Dr. James H. Finke (Cleveland Clinic), and is regularly monitored for contaminations.

### Plasmids and site directed mutagenesis

Plasmids 3xFLAG-dCas9/pCMV-7.1 (Addgene plasmid no 47948, RRID: Addgene_47948) and gRNA Cloning Vector Bbs-I ver 2.0 was obtained from Addgene (plasmid no 85586 RRID:Addgene_85586). T4 polynucleotide kinase (New England Biolabs, Cat. M0201S) was used to anneal and phosphorylate oligonucleotides 5’-CACCGCCTACAAGCTCAGAACGAGC-3’ and 5’-AAACGCTCGTTCTGAGCTTGTAGG C-3’ to make a double stranded DNA fragment. This DNA fragment was cloned into gRNA Cloning plasmid Bbs-I ver 2.0 (Addgene plasmid no. 85586, RRID:Addgene_85586) for targeting *GM2-synthase* promoter using enChIP method. pCT310 used for expression of 6X His-dCas9-3X FLAG in bacterial cells was obtained from Addgene (Plasmid no 111140, RRID:Addgene_111140). LPCX-SMAD2 (Plasmid No 12636 RRID:Addgene_12636), pCMV5B FLAG-SMAD3 (Plasmid No 11742, RRID:Addgene_11742), pcDNA FLAG-

SMAD4M (Plasmid No 14959, RRID:Addgene_14959) were all obtained from Addgene to perform the over-expression experiments. To clone the Region A (-333 to +37) of *GM2-synthase* promoter into pGL3-Basic (Promega, Madison, WI, Cat: PR- E1751, Addgene_212936), first primers listed in Table S1 were used to amplify Region A (-333 to +37) from genomic DNA. The amplified DNA was then cloned into pTZ57R/T (Fermentas, Lithuania, Cat: K1214A) using T/A cloning method followed by subsequent subcloning into pGL3Basic using KpnI and HindIII (New England Biolabs, Beverly, MA, Cat: NEB #R3142 and R0104S). Deletions and mutations of SP1 binding site and SMAD binding elements were made using site directed mutagenesis with primers indicated in Table S1. pcDNA3.1-P300 was obtained from Addgene (plasmid no 23252, RRID:Addgene_23252). For shRNA mediated knockdown of p300 in HEK293T cells, Tet-pLKO-neo was obtained from Addgene (Plasmid No 21916, RRID:Addgene_21916) and Oligos 5’-CCGGCCAGCCTCAAACTACAATAAACTCGAGTTTATTGTAGTTTGAGGCTGGTTTTTG-3’ and 5’-AATTCAAAAACCAGCCTCAAACTACAATAAACTCGAGTTTATTGTAGTTTGAGGCTGG-3’ were annealed and cloned following digestion of Tet-pLKO-neo with AgeI and EcoRI. All constructs were verified by sequencing.

### enChIP-Real-Time PCR

enChIP was performed using protocol as described previously (19). Two sets 2×10^6^ HEK293T cells seeded in 100 mm culture plates were co-transfected with either 3xFLAG-dCas9/pCMV-7.1 and gRNA Cloning Vector Bbs-I ver 2.0 (Empty) or gRNA Cloning Vector Bbs-I ver 2.0 (Guide RNA). The cells were replated in 150 mm culture plates after 24 hrs of transfection, and were harvested and crosslinked using 37% formaldehyde in 37°C for 10 mins, 48 hrs following transfection. Chromatin fragmentation was done using Cole Palmer CPX500 ultrasonic probe sonicator with median fragment length 500bp (20). The sheared chromatin was then put through a pre-clearing step with 3 µg of mouse IgG isotype control (CAT: sc-2025, RRID:AB_737182) coupled with Dynabeads protein G (10004D) followed by overnight incubation with Dynabeads protein G conjugated to Anti FLAG antibody (Sigma CAT: F1804, RRID:AB262044). Following overnight incubation, the Dynabeads were washed for two times with wash buffer containing low salt (20 mM Tris, pH 8.0, 150 mM NaCl, 2 mM EDTA,1% TritonX-100, 0.1% SDS), wash buffer containing high salt (20 mM Tris, pH 8.0, 500 mM NaCl, 2 mM EDTA, 1% TritonX-100, 0.1% SDS), wash buffer containing lithium Chloride (10 mM Tris, pH 8.0, 250 mM LiCl, 0.5% IGEPAL-CA630, 1 mM EDTA, 0.5% sodium deoxycholate) and TBS (50 mM Tris-HCL p^H^-7.5,150 mM NaCl, 0.1% IGEPAL CA-630). The chromatin precipitated by the Dynabeads were then eluted with elution buffer having the composition of 50 mM Tris-HCL p^H^-7.5,150 mM NaCl, 0.1% IGEPAL CA-630 and 5 mg/ml 3X FLAG peptide (Sigma Aldrich F4799). The eluted chromatin was then incubated at 65°C overnight with 200 mM NaCl for reverse crosslinking. Following this, the protein component of the immunoprecipitated chromatin was digested using proteinase K for 2 hrs at 45°C, and the remaining DNA component was then purified. The DNA isolated through enChIP was analysed by applied biosystems 7500 Fast Realtime PCR using primers targeted against region A of *GM2-synthase* promoter as listed in Table S1. The predicted structure of dCas9 – guide RNA-target DNA complex was predicted using AlphaFold 3 (21) and visualised with ChimeraX (22) (RRID:SCR_015872). The protein sequence of dCas9 was obtained from Expasy translate tool (RRID:SCR_024703) using the coding sequence obtained from dCas9 expression vector used (Addgene plasmid no 47948, RRID:Addgene_47948). The protein sequence obtained from Expasy translate tool, the guideRNA sequence and the target DNA sequences were given as inputs in AlphaFold 3 (RRID:SCR_025885) and the output predicted structure was exported in PDB (RRID:SCR_012820) format and visualised in ChimeraX software.

### enChIP-Mass Spectrometry

enChIP-MS was performed with chromatin from 4×10^7^ HEK 293T cells immunoprecipitated with 30 µg of Anti-FLAG Antibody (Sigma CAT: F1804, RRID:AB262044) and 300 µl of Dynabeads protein G per condition following the same protocols as described for enChIP-real-time PCR. Precipitation of eluted samples were done using 1ml isopropanol, 50 µl 3M sodium acetate and 5 µl of 20 mg/ml glycogen at -20°C overnight. The immunoprecipitated samples were boiled with 2x loading dye for 30min before subjecting to polyacrylamide gel electrophoresis (PAGE). Thirty samples were taken from the gel for the protein digestion. The resulting bands were rinsed and destained using 50% ethanol and 5% acetic acid followed by drying with acetonitrile. Before the in-gel digestion, samples were reduced using DTT and alkylated with iodoacetamide. Every band was completely digested with trypsin in-gel by adding 10 μl of the 5 ng/μl of the enzyme to 50 mM ammonium bicarbonate following overnight incubation at room temperature. After formation of the peptides, the polyacrylamide was extracted using two aliquots of 30 μl of 50% acetonitrile mixed with 5% formic acid. Together, the volume of these extracts was reduced to about 10 μl in a speed-vac. The peptides were reconstituted with 0.1% formic acid to yield a volume of around 30 μl for LC-MS analysis, which was performed in the Proteomics core of the Lerner Research Institute of the Cleveland Clinic, Cleveland, OH, USA. The LC-MS system consisted of a Captive-Spray ion source and a Bruker Times-Tof Pro2Q-Tof mass spectrometry system, both from Bruker Daltonik, Bremen, Germany in positive ion mode. The reverse phase capillary chromatography column used in the HPLC analysis was a Bruker 15cmX75µm (i.d.) C18 Reprosil AQ, 1.9 µm, 120 Å. A single microliter of the extract from the column was added to the mass spectrometer’s online source using a gradient of acetonitrile and 0.1% formic acid at a flow rate of 0.3 microliters per minute. A Parallel Accumulation-Serial Fragmentation approach was employed to select precursor ions for fragmentation with a TIMS-MS scan and ten PASEF MS/MS scans, respectively, in the digest analysis. With a ramp time of 166 ms, the TIMS-MS survey scan was obtained between 0.60 and 1.6 Vs/cm^2^ and 100-1,700 m/z. The PASEF scans took 1.2 seconds to complete in total, and the collision energy used in the MS/MS studies ranged from 20eV (0.6 Vs.cm^2^). With the target value set to 20,000 a.u and the intensity threshold set to 2,500 a.u, precursors with two to five charges were chosen. For 0.4 seconds, precursors were dynamically excluded.

PEAKSOnline v11 was used to search the LC-MS/MS data using the human SwissProtKB database downloaded on 3-23-2022 (26,576 entries). Full tryptic peptides with no more than 2 missed cleavage sites were considered to perform these searches. The MS1 and MS2 mass accuracies were set at 20ppm and 0.06 Da respectively, carbamidomethylating was considered as a fixed modification, and oxidation of methionine and protein acetylation were considered as variable modifications. A reverse decoy database strategy was used to filter the results, Peptide, and protein FDR rates were set to 1%. A minimum of 2 peptides with at least one of these being a unique peptide identification were required to be identified for positively identified proteins. The mass spectrometry proteomics data can be accessed with the dataset identifier PXD065551 at the ProteomeXchange Consortium of the PRIDE (23) (RRID:SCR_003411) partner repository.

### Gene ontology, Pathway enrichment and CORUM complex analysis

The Gene Ontology analysis was done using Shiny GO Gene Ontology (24) webtool using GO-MF, GO-CCO and GO-KEGG functions with FDR cutoff set at 0.05 and using number of genes for ranking the pathways. The chord plot showing connections between all proteins and its corresponding pathways was generated with SR plots (25) using the GO Chord function. The CORUM pathway analysis (26) and the network clustering of all the identified complexes was done in Cytoscape (27) (RRID:SCR_003032) from the enrichment data generated from gProfiler (28) (RRID:SCR_006809). EnrichmentMAP (29) (RRID:SCR_016052) and AutoAnnotate (30) Cytoscape plugins were used to functionally cluster the network in its represented form.

### CLASP

CLASP assay in SK-RC-45 cells was done using modifications of method as described earlier (31). Briefly, 2×10^6^ SK-RC-45 cells plated in 15 cm tissue culture plates were fixed with 1% formaldehyde. Cells were then lysed and fragmentation of the chromatin with median fragment length 500bp was achieved using Cole Palmer CPX500 ultrasonic probe sonicator. The fragmented chromatin was cleared with hard centrifugation and diluted with NMNT buffer (10 mM Tris pH 7.9, 500 mM NaCl, 5 mM MgCl2, 0.05% Nonidet P-40). RNA-Protein (dCas9) complexation was achieved through incubating either only 0.16 mg purified dCas9-FLAG, or 0.16 mg of purified dCas9-FLAG complexed with an in-vitro synthesized guide RNA (G.S) targeting the region A of *GM2-synthase* promoter. The cleared fragmented chromatin was then subjected to overnight incubation with the gRNA-dCas9 complex. Following this overnight incubation, 50 µl M2 agarose resin (CAT: A36801) was introduced in the reaction to immunoprecipitate the chromatin bound to dCas9-guide RNA complex. After an incubation of 4 hrs., the M2 agarose resin was centrifuged and washed two times with 500 μl NMNT buffer before the subsequent elution of the RNP (ribonucleoprotein) bound chromatin complex using 0.32 mg/ml 3X FLAG peptide (CAT: SAE0194) in 0.1M NaCl NMNT buffer for 1 hrs at 37°C with shaking. The eluted chromatin was then incubated at 65°C overnight with 200 mM NaCl for reverse crosslinking, following which the protein component was digested using proteinase K (New England Biolabs, CAT: P8107S) for 2 hrs. at 45°C. The remaining DNA component was then purified and the isolated DNA was then analyzed by Applied Biosystems 7500 Fast real-time PCR (RRID:SCR_018051) using primers targeted against region A of *GM2-synthase* promoter as listed in Table S1. Identification of the proteins co -immunoprecipitated with the locus by the CLASP method was done by western blotting as described by *Di Giorgio et al* (32). Purification of dCas9 and *in -vitro* synthesis of designed guide RNA was performed according to *Tsui et. al* (31). For *in -vitro* synthesis of designed guide RNA, the forward primer was designed with custom guide sequence after a T7 RNA promoter which was subsequently annealed with a universal reverse primer for an overhang extension PCR step to make a ds DNA template for synthesis of the guide RNA. In the next step the dsDNA template was used with Hiscribe T7 RNA polymerase kit (NEB Cat no E2040S) to generate the synthesized guide RNA. This synthesized guide RNA was run on a 7M Urea PAGE after which the guide RNA band was excised and eluted for downstream experiments.

### Luciferase assay

Wild type, deletion or mutation variants of pGL3Basic Region A (-333/+37) were transfected to HEK-293T cells as depicted in Fig. 6. Cell harvestation was done 2 days post transfection and harvested cells were washed for two times with phosphate buffered saline (PBS) before their subsequent lysis with luciferase lysis buffer (Promega, Madison, WI). Following lysis, the cell lysate was cleared with a short spin at 6000 rpm and protein concentration was determined with BCA protein assay (Thermo Fisher scientific). 20 µl of equal protein dilutions were incubated with 30 µl of luciferase assay reagent (21) followed by Stop and Glo buffer (provided with luciferase assay kit). Varioscan Flash multimode reader (Thermo Fisher Scientific) was used to measure the luminescence.

### RNA isolation, cDNA synthesis and Realtime PCR

TRIzol reagent (Invitrogen) and Verso cDNA synthesis kit (Thermo Fisher Scientific) was used to extract total RNA from cells and to synthesise cDNA from RNA following extraction respectively. Power up SYBR master mix (Invitrogen, Cat: A25742) was used to perform Real-time PCR in a 7500 Fast real-time PCR system (Applied Biosystems). All primers were procured from integrated DNA Technologies which are listed in Table S1.

### ChIP assay

SK-RC-45 cells seeded in 10 cm tissue culture plates were fixed using 1% formaldehyde followed by treatment with 1mM glycine in PBS. The fixed cells were washed for two times with cold PBS, scraped and lysed with 1ml NP-40 lysis buffer (50 mM Tris HCl, pH 8, 10 mM EDTA, 1% SDS). The lysate was then centrifuged in 2000 rpm for 5 mins to collect nuclear pellet which was then washed for two times, pelleted and resuspended in 800µl MNase (micrococcal nuclease) digestion buffer (10 mM Tris HCl, pH 7.5, 15 mM NaCl, 60 mM KCl, 0.15 mM spermidine). Cell suspension was then incubated with 1mM CaCl_2_ at 37°C for 5 mins, following which the chromatin was fragmented with 30 units MNase for 10 mins at 37°C. The fragmentation was terminated with 200 µl 5X nuclear lysis buffer (50 mM Tris HCl, pH 8, 10 mM EDTA, 1% SDS) and the lysate was cleared by a hard spin of 10,000 rpm for 10 mins at room temperature. The supernatant from this step was then collected and 10 times diluted with ChIP dilution buffer (1.1% Triton X-100, 1.2 mM EDTA, 16.7 mM Tris-HCl, pH 8, 167 mM NaCl). The chromatin was allowed to react with appropriate antibodies against either Sp1 (CAT: D4C3, 9389, RRID:AB_11220235), SMAD 2/3 (CAT: 5678S, RRID: AB_10693547) or SMAD 4 (CAT: 38454S, RRID: AB_2728776) from Cell Signalling Technologies (CST) and immunoprecipitated with Dynabeads protein A/G beads (Invitrogen). Thereafter, immunoprecipitated beads were washed twice each with high salt wash buffer, low salt wash buffer, lithium chloride wash buffer and TE, and the chromatin was released from the beads with elution buffer and was reverse crosslinked with 200mM NaCl at 65°C overnight, following which, the samples were digested with proteinase K and DNA was purified.

### Immunoprecipitation, sub-cellular fractionation and immunoblot analysis

10 cm culture plates seeded with cells were harvested by scraping and incubated with IP buffer (50 mM Tris-HCL pH-8, 10 mM KCL, 1 mM EDTA, 0.5% NP-40, 10% Glycerol) and nuclear extraction buffer (20 mM HEPES p^H^-7.9, 10 mM KCL, 1 mM EDTA, 20% Glycerol, 400 mM NaCl, 0.2% PMSF). Following lysis, cell debris were removed with centrifugation at 12000 rpm at 4°C for 30 mins. Equal amount of total protein was then incubated with SP1 antibody (CAT: D4C3, 9389, RRID:AB_11220235), SMAD 2/3 (CAT: 5678S, RRID: AB_10693547) or SMAD 4 (CAT: 38454S, RRID: AB_2728776) overnight, following which immunoprecipitation was achieved using Dynabeads protein A or G for 4 hrs at 4°C. Immunoprecipitated complexes were washed twice with nuclear extraction buffer, boiled for 10 mins with 2X loading dye and subjected to western immunoblotting. Cytosolic fractions were obtained by cell lysis with hypotonic buffer solution (20 mM Tris –HCL p^H^-7.4, 10 mM NaCl, 3 mM MgCl_2_). The supernatant was collected as cytoplasmic fraction after centrifugation at 3000 rpm for 10 mins at 4°C and the pellet containing the nucleus was washed twice with hypotonic buffer solution before being lysed with cell extraction buffer (10 mM Tris-HCL p^H^7.4, 2 mM Na_3_VO_4_, 100 mM NaCl,1%Triton-X100, 1 mM EDTA,10% glycerol, 1 mM EGTA, 0.1% SDS, 1 mM NAF, 0.5% sodium deoxycholate, 20 mM Na_4_P_2_O_7_). Western blot was done as described previously (2). As loading control of nuclear fraction, Lamin A/C (PA5-116716) levels were monitored.

### *In-silico* computational modelling

The three-dimensional structure of SP1 was obtained from the AlphaFold Protein Structure Database (33, 34), which web-resource is known to reliably predict models of proteins in an all-atom description. Since there was no experimentally resolved structure of the full SMAD trimer available, except a trimer of its core region (PDB: 1U7V), the full structure of the trimer SMAD was modelled using AlphaFold webserver (34). Three different compositions of the complexes were modelled, as the following: DNSPAD (for DNA, SP1 & SMAD complex), DNSP (DNA+SP1 complex) and DNAD (DNA+SMAD complex). These all-atom models were then processed in the CHARMM-GUI web server to set them up for further calculations (35). A basic level of refinement of each model was done using CHARMM36m (36) force field-based energy minimization, using Steepest Descent (23) and Adopted Basis Newton-Raphson (ABNR) energy minimization algorithms sequentially, while applying slight restraints on the atoms. Energy minimization was done in vacuum and in implicit solvent environment using GBSW model (37).

### Immunostaining

SK-RC-45 cells were stained with SMAD 4 (38454S, RRID: AB_2728776) and SMAD 2/3 (CAT: 5678S, RRID: AB_10693547) and hamster monoclonal anti-human GM2 antibody (DMF10.167.4, a gift from Dr. Kenneth Rock, University of Massachusetts Medical School, Worcester, MA and Dr. James H. Finke, Cleveland Clinic, Cleveland, OH) antibodies by methods described previously (2, 12). Glass cover slips seeded with SK-RC-45 cells were treated with sodium butyrate for indicated amount of time. Post treatment, 3.7% formaldehyde was used to fix the cells and were subsequently permeabilized using 0.1% Triton X-100 in 1X PBS. The cells were then incubated with respective primary antibodies with 1:50 dilution overnight. Following morning, the cover slips were washed thrice with 1X PBS before incubation with Alexa fluor -488/595 goat anti hamster antibody (CAT: A-21110, RRID:AB_2535759) and goat anti rabbit antibody Alexa fluor 488 (CAT-A11008,RRID:AB_143165). The nucleus of the cells was stained with DAPI.

### Statistical Analysis

Student’s t-test were used to calculate the p-values in GraphPad Prism (RRID:SCR_002798) software. In each case statistical significance was considered with (*) *p<0.05, (**) p<0.01 and (***) p < 0.001*.

## Results

### Targeting dCas9 to the promoter region of *GM2-synthase* gene enabled successful enrichment of the desired locus

The transcriptional regulation of the *GM2-synthase* gene is complex and not yet fully understood. Our previous results reflect an exquisite mechanism involving the transcription factor SP1 which recruits the repressor, HDAC1 at the *GM2-synthase* TSS thereby repressing its transcriptional outcome in normal kidney epithelial cells or in conditions that repress *GM2-synthase* transcription. This SP1-HDAC1 mediated repression is reversed in renal cancer cells (which over-express *GM2-synthase*) through an acetylation-dependent proteasome mediated degradation of the repressor, SP1 as discussed in *Banerjee, A. et. al., 2019* (18). Mechanistically, although withdrawal of the repression owing to SP1 degradation contributes towards the observed activation of *GM2-synthase* transcription, the fact that *GM2-synthase* transcriptional activation may be through a distinct mechanism other than the de-repression cannot be ruled out. This demands identification of the DNA- as well as non-DNA binding partners involved in *GM2-synthase* transcription which necessitated a two-step profiling of the entire proteome associated with *GM2-synthase* promoter which comprised of two different dCas9 mediated proteome profiling tools (Fig. 1 & Fig. 4). To address this, we applied a previously reported proteomic profiling approach based on the dCas9 mediated enChIP technique as the first, hypothesis generation step of this two-step proteome profiling regime (19). Fig. 1A depicts the entire enChIP method in a schematic form. For the enChIP-MS method, first a catalytically dead Cas9 molecule (expressed from 3xFLAG-dCas9/pCMV-7.1 following transfection) is directed to the locus of interest with the help of a specific guideRNA (expressed from gRNA Cloning Vector Bbs-I ver 2.0 cloned with guide RNA targeting *GM2-synthase* TSS following transfection). Following targeting of the dCas9 to the intended locus, the chromatin is crosslinked, sonicated to desired fragment length and immunoprecipitated with help of appropriate antibody to isolate the intended locus, bound with its locus specific proteome. The DNA component of this immunoprecipitated complex is then analysed with RT-PCR to ensure the specific enrichment of the intended locus (Fig. 1F) and the proteome is identified with Mass spectrometric analysis. For this work, the enChIP-MS method was utilised as a hypothesis generating engine, that acted as the first step of a two-step proteome profiling pipeline, in which it nominated players for identification through an even more sensitive second step CLASP-WB (*in-vitro* dCas9 mediated proteome profiling method) described in detail in the following segments (Fig. 4). In order to profile the chromatin associated regulatory elements present across a genomic region more than 500 bp long (Fig. 1B), the average length of the chromatin fragments were obtained at a median length of 500 bp with sonication (Fig. S1A). This particular region of chromatin which encompasses the *GM2-synthase* transcription start site (TSS) and spanned across a length of around 500 bp, was selected on the basis of our previous publication (18) which established the importance of the histone modifications and regulatory activity of this stretch of chromatin on *GM2-synthase* transcription. To profile the total regulatory elements associated with the mentioned genomic region, a FLAG tagged dCas9 molecule was directed to the locus with the help of a single guide RNA (sgRNA) designed against a genomic location upstream of the promoter (Fig. 1B). Fig. 1C provides a visual context of the exact genomic location where the complex comprising dCas9, the used guide RNA and the targeted region of *GM2-synthase* promoter is expected to be formed. Cells transfected with the FLAG-tagged dCas9 clearly shows over-expression of dCas9 using an anti-FLAG antibody indicating that the over-expression system is functional (Fig. 1D). 2×10^6^ HEK-293T cells were co transfected with plasmids expressing FLAG tagged dCas9 and sgRNA expression vector cloned with the designed sgRNA, following which nuclear extracts were subjected to immunoprecipitation with anti-FLAG antibody. This immunoprecipitation showed a robust enrichment of dCas9 when compared to immunoprecipitation with normal isotype control (Fig. 1E). This result depicted the efficiency of the anti-FLAG antibody in immunoprecipitating FLAG-dCas9, as depicted by the presence of the thick band in the IP condition in comparison to negligible band in the supernatant. High specificity of the IP process was also demonstrated by the absence of any protein in the isotype control. The enrichment of the specific genomic locus was then checked using enChIP real-time PCR (Fig. 1F) in which the ChIP assay was performed using anti-Flag antibodies with a targeting set expressing dCas9 along with a targeting guide RNA, and a non-targeting set expressing only dCas9 acting as a control. These results confirm that the targeted chromatin is successfully enriched using the enChIP method.

**Fig 1.**
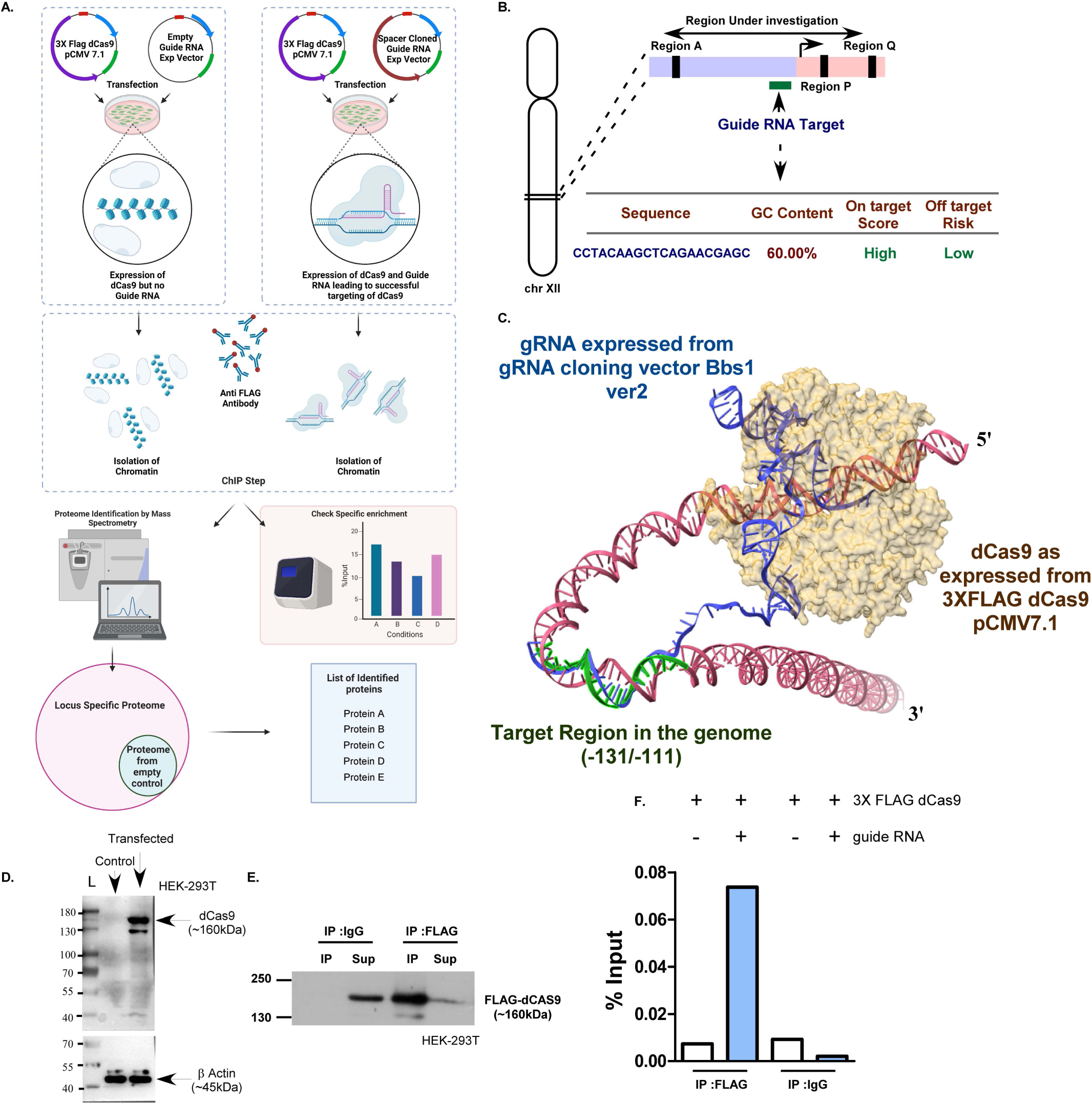
Efficient targeting and specific isolation of GM2 synthase locus through enChIP method. A. Schematic representation of enChIP-MS technique used as the first step of the two-stage proteome profiling regime. B. Genomic location of the target locus, intended target of the guide RNA, and sequence of the designed guide RNA. C. AlphaFold-predicted structural representation of dCas9-gRNA complex at target genomic locus. D. Expression of dCas9 after transfection of dCas9 expression vector in HEK-293T cells. E. Immunoprecipitation of FLAG tagged dCas9 with anti-FLAG antibody and isotype control, high relative enrichment of dCas9 in the FLAG-IP samples, depicts the efficiency of the immunoprecipitation. F. Enrichment of GM2 synthase locus with enChIP method. Data represents relative enrichment of the sample used for mass spectrometric analysis. Conditions 1-2 and 3-4 represent samples that were immunoprecipitated with specific anti -FLAG antibody and normal mouse IgG isotype control respectively. The blue bars represent targeting IP samples. Biological replicates showing efficiency and repeatability of dCas9 targeting to the intended locus is given in Fig S1.

### Engineered DNA binding molecule mediated chromatin immunoprecipitation (enChIP)-MS identifies critical, robust and diverse group of proteins associated with the *GM2 - synthase* locus

The enChIP-MS analysis identified 30 proteins exclusively present in the non-targeting sample (dCas9 only), 1757 proteins exclusive to the targeting sample (dCas9 + Guide RNA) and 2170 proteins significantly enriched in the targeting (dCas9 + Guide RNA) sample over its non-targeting (dCas9 Only) counterpart (Fig. 2A). To exclude non specificity, all the proteins which were either present exclusively in the targeting sample or were having a significant high enrichment compared to the two internal controls were selected (Fig. 2B) and were subjected to sequential sorting using gene ontology (10) enrichment analysis based on their molecular functions (MF), cellular component (CC) and KEGG pathway analysis in the mentioned order (Fig. 2C). This GO-MF enrichment analysis revealed that the predominant proteome identified is associated with molecular functions related to nucleic acid binding (Fig. 2D). Further, this finding not only provided us with a large pool of proteins identifiable as regulators of *GM2-synthase* transcription, but also highlighted the efficiency of the enChIP-MS system to target dCas9 to the intended locus with a low extra-chromosomal background. With this step allowing us to focus on only the “nucleic acid binding” proteins identified through enChIP-MS, these nucleic acid binding proteins were then subjected to gene ontology (10) enrichment analysis on the basis of cellular components (10). This analysis further segregated them and revealed nucleic acid binding proteins filtered and selected in the previous step, to be a versatile group associated with many cellular locations such as the nuclear lumen as well as chromosome and chromatin (Fig. 2E). This approach to narrow down on the proteins responsible for *GM2-synthase* transcriptional regulation, from a large group of functionally diverse proteins that were identified through MS, now provided us with a pool to search for the individual transcriptional regulators with the intricate roles that each of them may play in the regulation of *GM2-synthase* transcription. The proteins enriched in “Chromosome and Chromatin” GO Cellular component step (Fig. 2E) were subjected to KEGG pathway enrichment analysis (Fig. 2F), which showed most of the identified proteins enriched with chromosome and chromatin were also associated with pathways critical to cancer, such as “Transcriptional mis-regulation in cancer”, “Viral carcinogenesis” or “microRNAs in cancer” in addition to classical chromosomal maintenance pathways like “DNA replication”, “Homologous recombination” pathway or pathways related to “Cell cycle” regulation. Fig. 2G depicts the network of each individual protein with the pathways that they were enriched in.

**Fig 2.**
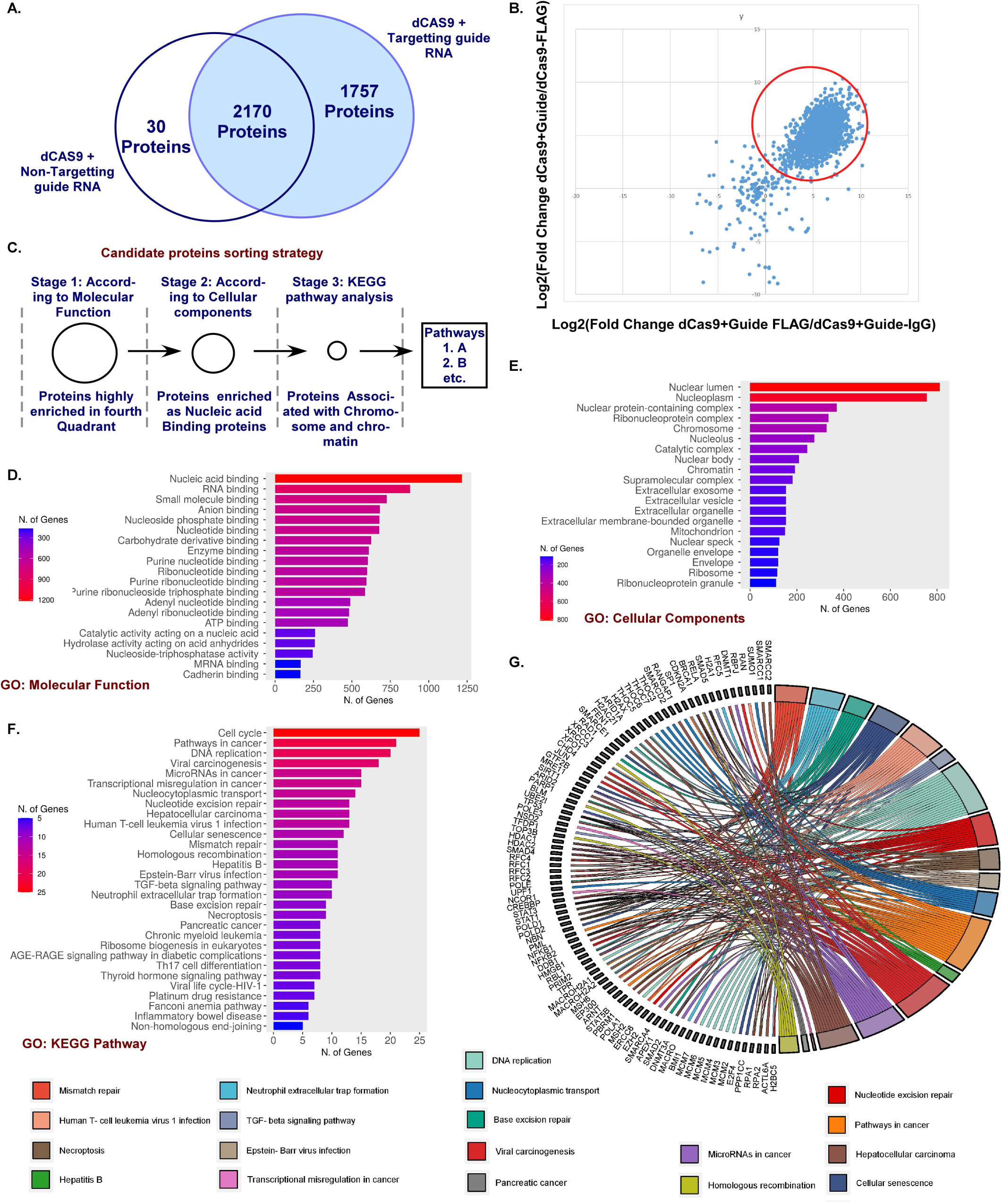
Targeted promoter capture identifies a robust and diverse proteome associated with GM2 Synthase promoter in the first step. A. Schematic representation of all the proteins identified through enChIP-MS technique. White circle on the left is nonspecific control and the blue circle in the right is the specific sample containing both dCas9 and target guideRNA, while the proteins shared in both have a significantly high enrichment in the specific sample (LFQ ratio ≥20). B. Scatter plot showing relative enrichment of the identified proteins with respect to both specific-nonspecific and targeting-non targeting internal controls. C. Strategy for sorting candidate proteins to nominate most prominent candidates for the second CLASP-WB step. D. Categorization of all identified proteins with respect to their Gene Ontologies – Molecular Function. E. Categorization of the nucleic acid binding proteins with respect to their cellular location. F&G. Categorization of the Chromosome and chromatin associated proteins with respect to KEGG pathway enrichment and depiction of each of the chromosome and chromatin associated proteins to their corresponding pathways as assigned by KEGG.

### CORUM Complex analysis identifies protein complexes important for transcriptional regulation from large pool of enChIP identified chromatin proteome

The proteins enriched in chromosome and chromatin component (Fig. 2E) were further subjected to CORUM (Comprehensive Resource of Mammalian Protein Complexes) protein complex enrichment analysis (26), which displayed proteins known to form significant protein complexes, many of which are well reported to be associated with transcriptional regulation in cancer. Fig. 3A depicts a network analysis of all the identified protein complexes identified as chromatin and chromosome associated protein in enChIP MS technique. Here the nodes represent the individual protein complexes, the clusters are larger functional groups, while the edges represent the relation between the individual protein complexes. From this large list of complexes, protein complexes associated with the previously reported regulators (Such as SP1 and HDAC1) were selected. Fig. 3B depicts the important complexes identified through CORUM complex enrichment analysis as well as the constituent subunits of those complexes which were present in our enChIP-MS analysis. This complex enrichment analysis revealed the plausible association of SMAD2/4 containing complexes as well as the presence of p300 in complexes having both SP1 and SMAD 2/4 as their components. Interestingly, p300 is well reported to modify and regulate both SP1 (38) and SMAD2/4 complex (39), which provided us with a basis to look further into the possibility of p300 and SMAD 2/4 complexes as regulators of *GM2-synthase* transcription. PPI interaction analysis in STRING-DB of all the components of these complexes (Fig. 3C) also showed a high confidence for the association between SP1-HDAC1, SMAD 2/4 and p300 (Fig. 3C) which further strengthened our hypothesis of SMAD2/4 and p300 as potential regulators of *GM2-synthase* transcription. Together, as the first step, enChIP-MS generated the hypothesis *de-novo,* and nominated mainly transcription factors and transcription factor associated co regulators as the best candidate pool to specify the important players to be validated and further investigated in the second more sensitive dCas9 mediated CLASP-WB step.

**Fig 3.**
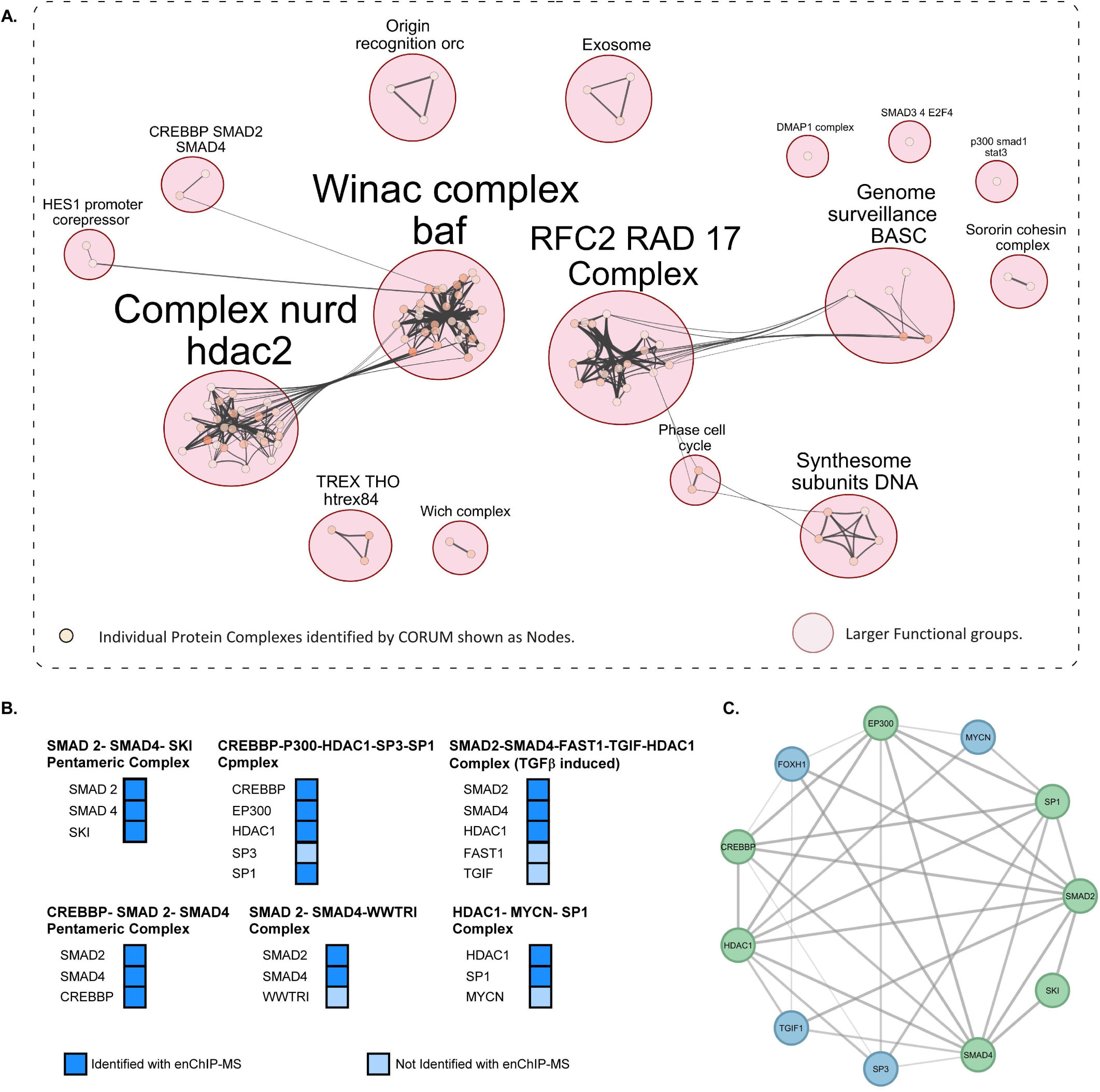
CORUM Complex analysis identifies important regulators from enChIP identified chromosome and chromatin associated proteins. A. Network analysis of chromosome and chromatin associated proteins through CORUM. Networking is represented as determined by gProfiler (p□0.05). Nodes represent protein complexes that were clustered into network by EnrichmentMAP. Autoannotate plugin was used to assign larger functional clusters to the network of nodes. B. Important CORUM Complexes and their components on the basis of their identification with enchIP. Dark blue boxes represent components that were identified in enChIP- MS method, while the light blue boxes represent the protein components that were not. C. STRING interactome of all the protein components of the selected CORUM complexes. The proteins marked in green represent the components identified by enChIP MS and the blue nodes represents the proteins that were not. The edge thickness represents the string interaction scores. (String edge cutoff was set at 0.45).

### CLASP-WB confirms the presence of important identified proteins in *GM2-synthase* TSS

As the second step of the two-step proteome profiling regime employed to decipher the regulatory mechanism at the *GM2-synthase* locus, presence of the candidate proteins nominated from the enChIP step was validated in a low *GM2 -synthase* expressing renal cancer cell line SK-RC-45. For this second step, serving as the initial platform to a robust array of orthogonal validations, we took help of a principally similar, yet a slightly different approach namely, CLASP (31). In this CLASP method an *in-vitro* synthesized guideRNA (Fig. S2H&I) and a purified dCas9 (from *E. coli*) (Fig. S2C&D) are utilized to constitute the locus targeting RNA-protein complex (Fig. 4A) while a non-targeting RNA-protein complex is used as a control (constituted with purified dCas9 and an *in-vitro* synthesized control guideRNA). Here, all the intermediate steps, both for synthesis of the guide RNA *in-vitro*, and for the purification of dCas9 are depicted in Fig. S2E-G and Fig. S2A-C respectively and are described in detail in the methods section. Here, the immunoprecipitation of the locus was done using the mentioned targeting and non-targeting RNA protein complexes with the chromatin isolated from SK-RC-45 cell line (Fig. 4A). Fig. 4B depicts the enrichment of the locus of interest in comparison with non-targeting background, which clearly shows achievement of desired relative enrichment of the locus of interest with low background (dCas9+ target gRNA versus dCas9+ control gRNA). Following this, the important proteins shortlisted from the enChIP-MS step are identified through western blot post immunoprecipitation of the locus using CLASP method. Here Fig. 4C to 4J depicts the western blots showing the presence of proteins that co-immunoprecipitated with our locus of interest. The results clearly show SP1, SMAD 4, SMAD 2, HDAC1, STAT3, p300, p53, p65, c-JUN, TEAD-1, SMARCE1, to significantly coimmunoprecipitate with the desired locus of the chromatin in the targeting samples (dCas9+ Target gRNA) compared to the non-targeting samples (dCas9 + scrambled gRNA), while Fig. 4I shows the absence of the protein IKB α as a negative control, which repeatedly remained un-detected in the targeting as well as the non-targeting samples. This further validated the hypothesis generated with the first enChIP step and identified the key proteins at the *GM2-synthase* locus in a new renal cancer cell line system. Here, the presence of FLAG-tagged dCas9 was used as a loading control to validate successful and equal pulldown of the chromatin and all CLASP-WB validations were performed with at least three biological replicates resulting in consistent enrichment patterns.

**Fig 4.**
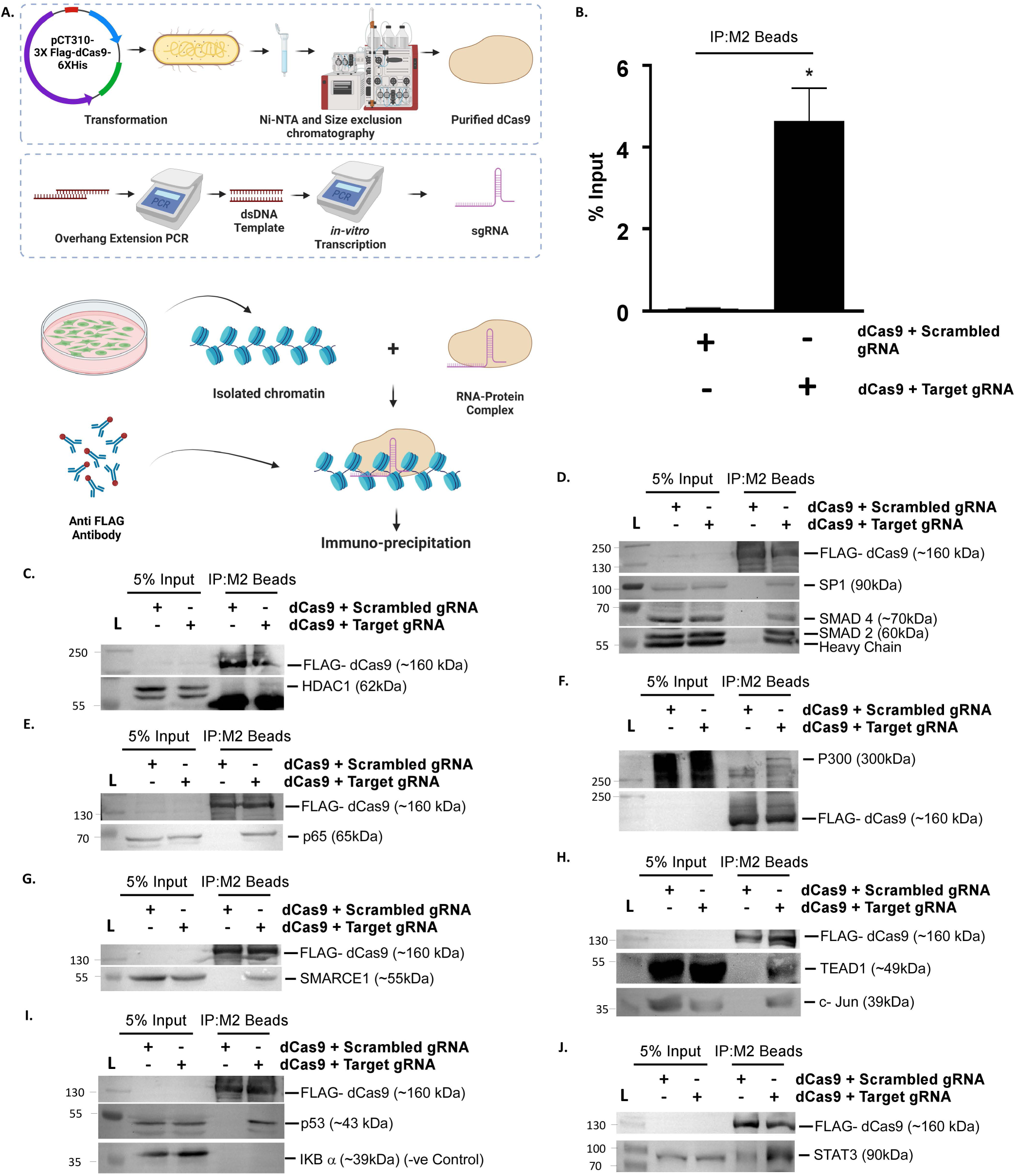
Second CLASP-WB step re-identifies important chromosome associated regulator proteins on GM2 synthase locus of SK-RC-45 cells. A. Schematic representation of CLASP technique. B. Relative enrichment of GM2 synthase locus with CLASP method. C-J. Identification of salient chromatin associated proteins co-purified with GM2 synthase promoter by CLASP-WB method from three different experimental repeats for each candidate. The other two repeats are represented in Fig. S5. Error bars represent mean ±S.E.M. of at least three independent determinations. (Student’s t test; *p 0.05**, p 0.01; ***, p 0.001). ns, not significant.

### SMAD binding element present upstream of *GM2-Synthase* TSS has an activatory role in *GM2-synthase* gene transcription

Following the identification of critical transcription factors as a major group of proteins in our two-step proteomic profiling pipeline (Fig. 2,3 & 4), we further looked for plausible transcriptional activators associated with the *GM2-synthase* promoter. SMAD 2 and SMAD 4 not only came up in the CORUM analysis of the MS data (Fig. 3B) as well as in CLASP-WB validations (Fig, 4), but also a presence of their binding site was predicted with TFind webtool at position -324/-318 (40–43) (Table. S2). To investigate the functional role if any, of the SMAD 2/4 complex in the transcriptional regulation of *GM2-synthase* gene, we treated a low *GM2-synthase* expressing renal cancer cell line SK-RC-45 with TGF β, a well-known positive modulator of SMAD 2/4 activity (44). In fact, TGF β treatment caused an expected time-dependent upregulation of p-SMAD 2 expression (Fig. 5F) which further correlated with an increase in the *GM2-synthase* message level (Fig. 5A), suggesting a plausible role of SMADs in *GM2-synthase* transcription. This was also supported by a significant increase in *GM2-synthase* mRNA expression upon SMAD 2/4 over-expression in HEK-293T cells (Fig. 5D). Interestingly, both TGF β (Fig. 5B & 5C) treatment and SMAD 2/4 over-expression (Fig. 5E & S3D) resulted in an even more amplified up-regulation of ganglioside GM2, the downstream metabolic product of *GM2-synthase* enzyme that is reported to be responsible for exerting pro tumorigenic effects (45, 46). This indicates that the metabolic output of ganglioside GM2 is highly sensitive to any changes in GM2 synthase transcript levels, and therefore to effects exerted by any of its transcriptional modulators. To confirm that the TGF β mediated activation of *GM2synthase* transcription involves the predicted SMAD binding element, the genomic region -333/+37 was cloned under a luciferase reporter, over-expressed in HEK-293T cells followed by either TGF β treatment, or over-expression of SMAD 2/3/4. A significant increase in the relative luciferase activity was observed in HEK cells upon treatment with either TGF β (Fig. 5H) or SMAD 2/3/4 over-expression (Fig. 5I), indicating a direct role of the SMAD-binding element in TGF β-induced *GM2-synthase* transcription. Further, to confirm the activation of TGF β/ SMAD signalling axis, changes in mRNA levels of canonical SMAD target gene Snail was monitored as positive control, which showed expected changes in response to both the inductions (Fig. S3A & S3B) as reported by earlier reports (47). Also, consistent with the previous reports, c-myc levels were also downregulated upon TGF β treatment (Fig. S3A) (48). Since, global HDAC inhibition was already shown to increase *GM2-synthase* transcriptional activity, we investigated the involvement of SMAD 2/4 complex upon HDAC inhibition by NaBu. NaBu resulted in a time- dependent increase in SMAD 2/3, as well as SMAD 4 association with the *GM2-synthase* TSS as demonstrated by ChIP assay (Fig. 5G). This is further supported by the fact that NaBu treatment in SK-RC-45 cells resulted in nuclear localisation of SMAD 2/4 complex (Fig. 5J) as shown by western immunoblot demonstrating a time-dependent increase in SMAD 2/4 in the nuclear, and a corresponding decrease in the cytoplasmic fraction. Nuclear localization of the SMAD 2/4 complex was also evident from immunofluorescent microscopy, which clearly showed a time-dependent localization of the SMAD 2/3 in the nucleus as evident from the co-localization of SMAD 2/3 with DAPI (Fig. 5K). Since, nuclear localization of SMAD 2/3 is triggered by its phosphorylation (49, 50), we checked for SMAD 2 phosphorylation. Data shows a time-dependent phosphorylation of SMAD 2 (Fig. 5L), indicating activation of Smad 2 (51). Collectively, these evidences suggest that SMAD 2/4 complex contributes to the regulation of *GM2-synthase* transcription. Notably, SMAD mediated transcriptional regulation and the amplitude of its effect on downstream target genes is regulated by multiple factors. For some genes like PAI-1and JUN B, the amplitude of SMADs effect is impacted by multiple binding sites (For example 3 SBEs in PAI-1 and 2 in case of JUNB)(1, 52), and in some genes the amplitude of the effect is determined by SMAD regulated enhancers and co-activators like in case of SNAI1(53). Although the induction of the single SMAD binding element present at position -324/-318 of *GM2-synthase* promoter showed induction of transcriptional activity proportionate to the over-expression of *GM2-synthase* mRNA in renal cancer cell lines (3-10 folds when compared to normal kidney epithelial cells) and in RCC tumours (3-10 folds when compared to tumour adjacent tissue)(18) along with an amplified induction of ganglioside GM2 (Fig, 5B,C,E & S3F) levels, involvement of co-regulators in other genes prompted us to investigate the roles of proximally present transcription factors on the inducibility of this SMAD binding element ultimately revealing an active antagonistic regulatory circuit.

**Fig 5.**
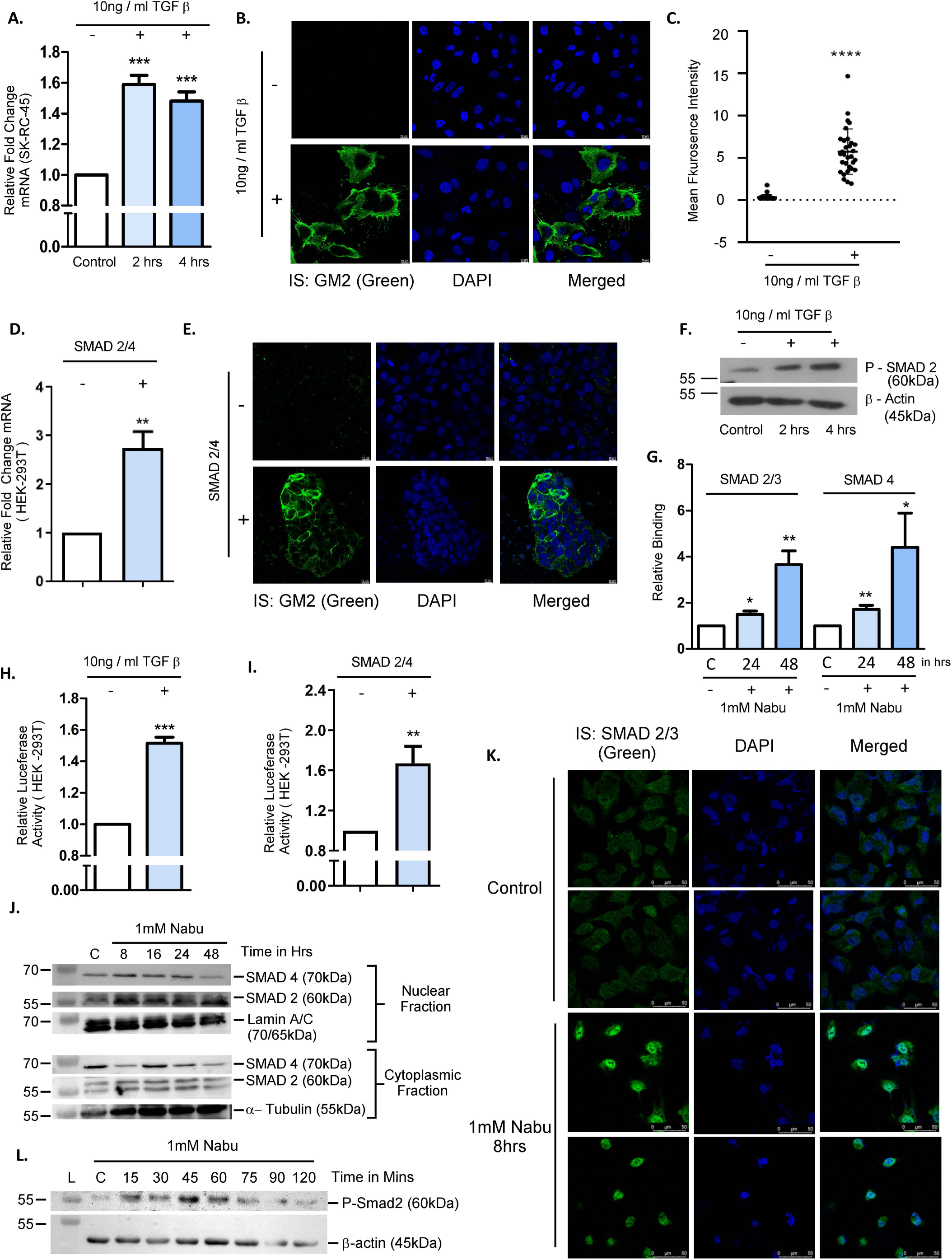
SMAD 2/4 Complex acts as a transcriptional activator of GM2 synthase gene. A. Relative fold change of GM2 synthase mRNA upon TGF β treatment. B&C. Change in ganglioside GM2 levels (Green) upon TGF β treatment and quantification of mean fluorescence intensity with respect to control cells. D&E. Relative fold change of GM2 synthase mRNA upon SMAD2/4 Over-expression and corresponding changes in ganglioside GM2 levels (Green) under the same condition. F. Induction of p-SMAD2 levels with TGF β treatment. G. Relative binding of SMAD2/4 complex with their predicted binding sites upon global HDAC inhibition. H&I. Luciferase promoter activity of GM2 synthase promoter upon TGF β treatment and SMAD2/4 over-expression. J&K. Nuclear localization of SMAD2/4 complex upon global HDAC inhibition. L. SMAD activation mark phosphorylation upon global HDAC inhibition. Error bars represent mean ±S.E.M. of at least three independent determinations. (Student’s t test; *p 0.05**, p 0.01; ***, p 0.001). ns, not significant. Fig 5F was performed twice with all other experiments of this figure performed at least three times. All statistical significance is calculated with respect to the individual untreated and empty vector control sets as shown in the individual sub-figures.

### Proximal SP1 binding site antagonizes the effect of SMAD2/4 binding element on *GM2 - synthase* transcription

In addition to the SP1 binding site (+109/+125) reported earlier (18), presence of another SP1 binding site proximal (-311/-295) of the TSS was predicted by TFind webtool (40, 54) (Table. S2). The presence of this upstream SP1 binding site only 6bp away from the earlier mentioned SMAD 2/4 binding element (SBE) along with the repeated identification of both SMAD 2/4 and SP1 in our two-step proteome profiling regime (Fig. 2, 3 and Fig. 4), prompted us to investigate the effect of this new SP1 binding site on *GM2-synthase* transcription, as well as the interplay, if any between the SP1 binding site and the SMAD 2/4 binding element under investigation. To address this, a small genomic region (-333/+37) harbouring the SP1 binding site and the SMAD 2/4 binding elements were cloned under a luciferase reporter gene. To understand the role played by this SP1 binding site in isolation from the rest of the regulatory players, the SP1 binding site under investigation was mutated at position +109/+125. This mutation resulted in an up regulation of relative luciferase activity (Fig. 6A) when compared with the wild-type condition, indicating that this SP1 binding site acts as a transcriptional repressor for *GM2-synthase* transcriptional activity. To further understand the effect of the SMAD 2/4 binding element, a 122 bp region encompassing both the SP1 binding site and the SMAD 2/4 binding element was deleted. Simultaneous deletion of both the SP1 as well as SBE resulted in a much lower relative luciferase activity in comparison to the SP1 mutated condition as well as the wild type condition, suggesting that the SMAD 2/4 binding element act as a positive modulator of *GM2-synthase* promoter activity (Fig. 6A). This observation was further supported by mutational studies in which the SMAD 2/4 binding element was mutated via site directed mutagenesis keeping the SP1 binding element intact, and vice-versa. Expectedly, this resulted either in a significant increase in the promoter activity upon SP1 mutation, or decrease upon SMAD 2/4 mutation (Fig. 6B). Interestingly, mutation of both the SP1 as well as the SMAD binding sites did not significantly alter *GM2-synthase* promoter activity compared to the wild-type, indicating again that anticipated activation owing to mutation-induced loss of SP1 binding is significantly compensated by the loss of promoter activity due to mutation-mediated loss of SMAD 2/4 binding. These results were further supported by experiments where we induced the system with all three inducers of *GM2-synthase* as shown in Fig. 5, namely either SMAD 2/4 over-expression, NaBu or TGF β treatment. Under all three inductions, significantly higher levels of luciferase activity were observed under conditions where the activity of SMAD 2/4 binding element were kept intact, while the activity of SP1 binding site was eliminated (Fig. 6C, Fig. 6D & Fig. 6E), indicating again that the positive activity of the SMAD 2/4 binding element on *GM2-synthase* transcription was more pronounced in the absence of the repressive activity owing to loss of SP1 binding. Hence, these results suggest that loss in SP1-mediated repression may not be the only mechanism behind *GM2-synthase* transcriptional activity observed, but that SMAD 2/4 binding does have a profound effect.

**Fig 6.**
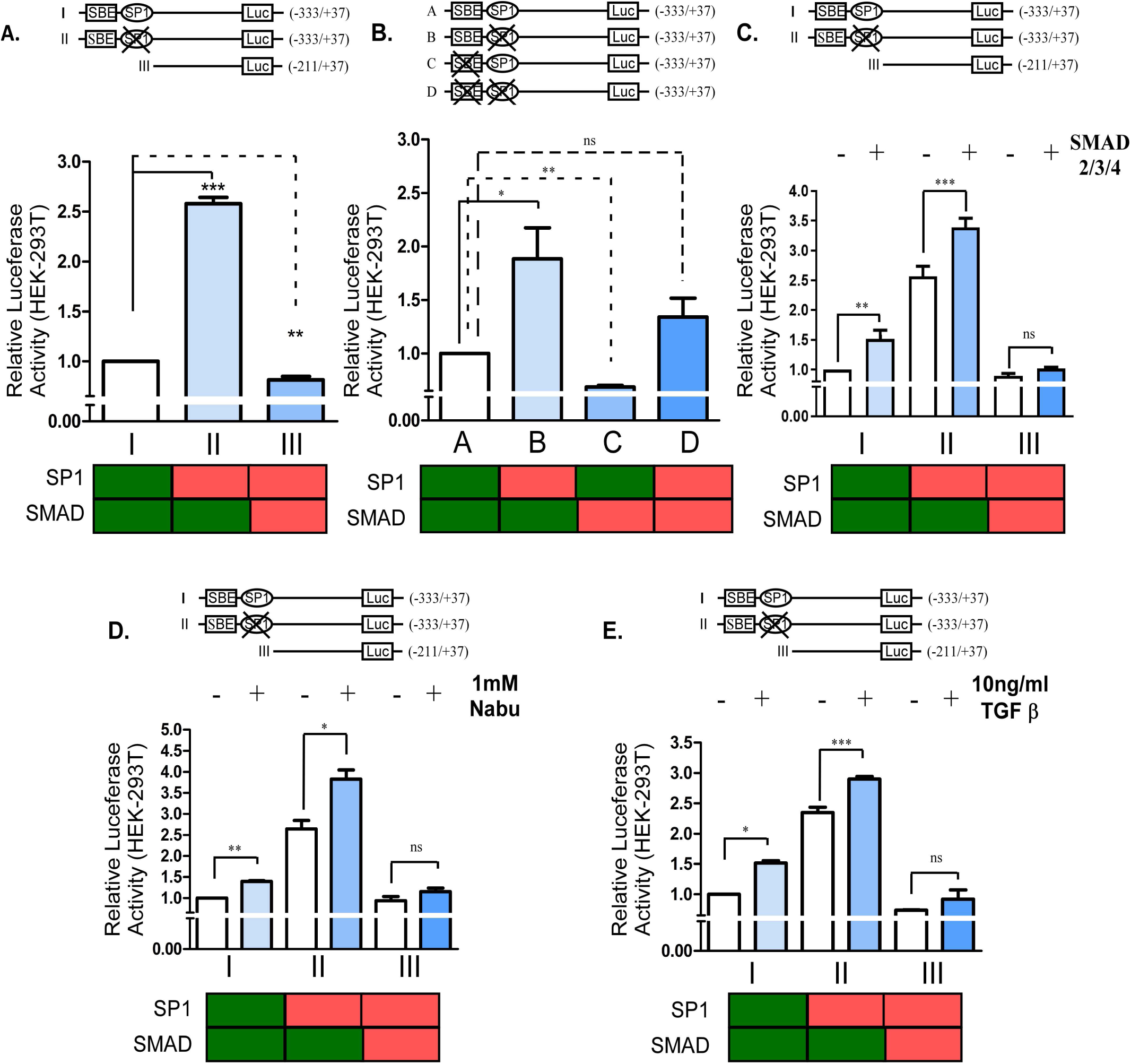
SP1 binding to the proximal SP1-binding site antagonizes the effect of immediate upstream SMAD 2/4 binding element in the transcriptional regulation of GM2 synthase gene. A&B. Dual luciferase assay involving deletion and mutation constructs showing the effect of proximal SP1 binding site and the SMAD2/4 binding element in GM2 synthase promoter activity C,D&E. Luciferase assay involving deletion constructs induced with Nabu, SMAD2/4 over-expression and TGF β treatment showing the inter-relation between the proximally present SP1 and SMAD2/4 binding elements in GM2 synthase promoter activity. Error bars represent mean ± S.E of at least three independent determinations. (Student’s t test; *p 0.05**, p 0.01; ***, p 0.001). ns, not significant.

### Computational modelling determines that simultaneous binding of SP1 and SMAD to their proximal binding sites is energetically unfavourable

In the relevant stretch of the DNA belonging to *GM2-synthase* promoter, the binding sites of SMAD and SP1 are only six nucleotides apart. Due to this close proximity of binding sites, given the large size of the proteins, the two proteins are likely to experience tremendous steric collisions if both of them simultaneously attempt to bind. Two models, DNSP (Comprising of SP1 bound to its DNA binding site) and DNAD (SMAD complex bound to SMAD binding element), were superimposed on each other by aligning them with respect to their common DNA segment, and the possibility of steric collision was very apparent, which has been presented in Fig. 7A. This model, although apparently hypothetical due to its impractical overlap in the model yet actually gives a plausible explanation why it was evidenced from the luciferase experiments (Fig. 6C, D and E) that the presence of SP1 binding site hinders the inducibility SMAD binding element. Often such constructed models quickly correct themselves to yield a relatively reasonable conformation by minimizing their internal strains due to steric repulsions, and so this possibility was tested by relaxing them through molecular mechanical force field-based energy minimization. The potential energy (PE) values of each complex in vacuum and under implicit solvent conditions have been tabulated in Fig. 7B. Since the model contained large number of atoms, to make the calculation faster and manageable, initially the minimization was done in vacuum, i.e. without adding the complexity of the solvent water. The results are tabulated in the Fig. 7B. The DNSPAD (Comprising of DNA, SP1 and SMAD) complex exhibited a total PE of −7803.1 kcal mol^-1^. In contrast, the DNSP and DNAD complexes showed PE of which indicates a destabilization of approximately 2046.5 kcal.mol^-1^ in favor of the separately bound binary complexes. Further, the energy minimization was done after adding the impact of solvation implicitly (using GBSW model), still avoiding the overload of the explicit solvent atoms. Under the implicit solvation conditions, the destabilization of the ternary complex compared to the two separate binary complexes was 10,549.5 kcal mol^-1^ (Fig. 7B) that gave stronger evidence of the impossibility of the ternary complex formation. None of these attempts of energy minimization actually led to any possibility of the stabilization of the larger complex.

**Fig 7.**
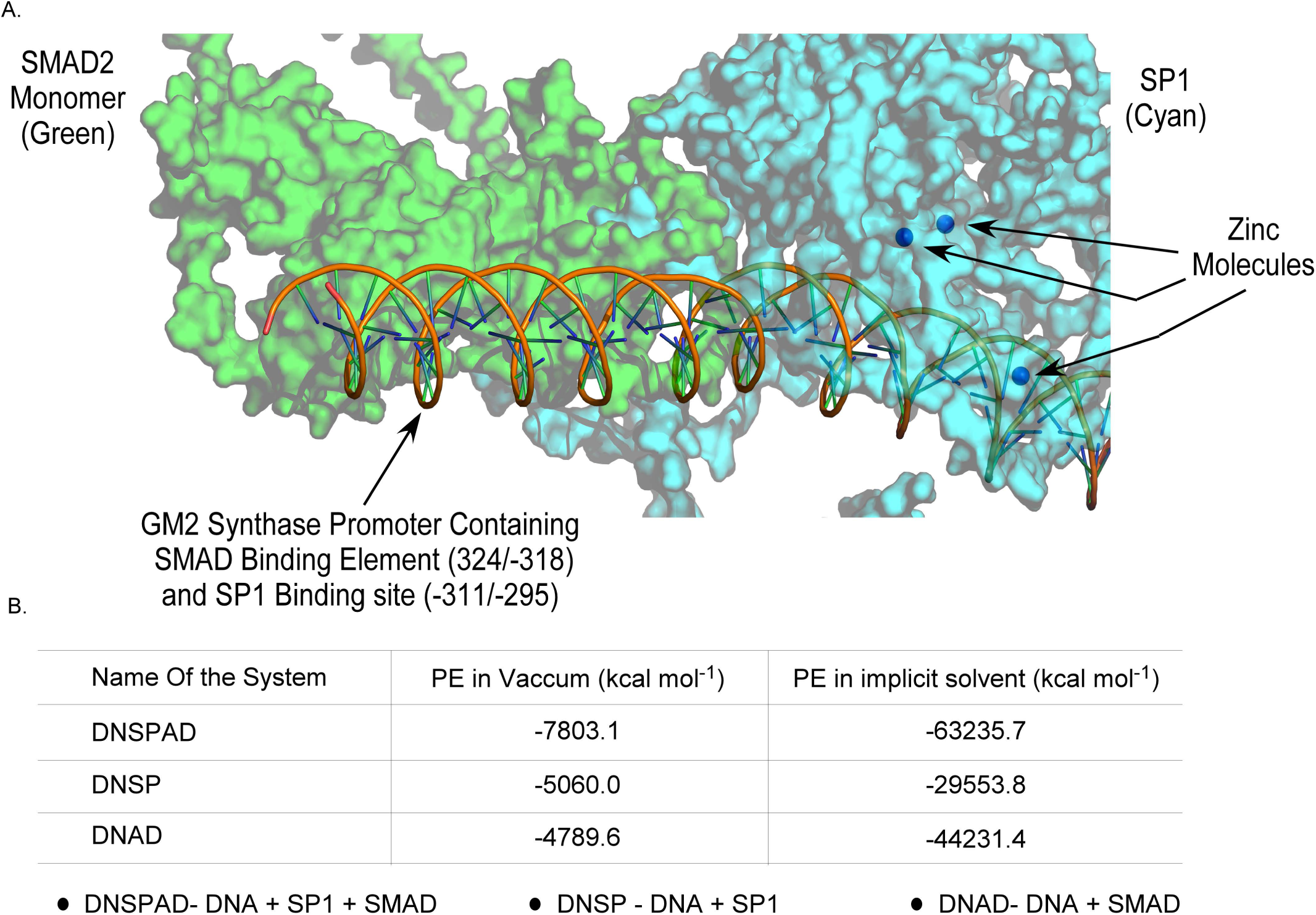
in-silico molecular docking results support SP1-SMAD2/4 functional antagonism for GM2 synthase gene. A. Surface representation of the SMAD2 monomer 1 (green) and SP1 (cyan) bound to DNA, illustrating spatial overlap indicating potential collision. Zinc atoms are shown as blue spheres, while the DNA double helix is depicted in orange using cartoon representation. Proteins are displayed in surface mode to highlight the interaction interface. B. Potential energy values (kcal mol^-1^) of different complexes calculated under vacuum and implicit solvent conditions.

Since the formation of the DNSPAD complex seems to be unrealistic, in this work there is no scope to take these models to any advanced level of calculations like molecular dynamics simulation-based sampling. Taken together, the computational modelling showed the simultaneous binding of SMAD and SP1 with their respective binding sites energetically less favorable due to steric constrains. This provided a possible explanation for the observed weakened impact of the SMAD binding element on *GM2-synthase* transcription, in presence of its proximal SP1 binding site.

### HAT p300 positively regulates *GM2-synthase* transcription

Since, acetylation-dependent degradation of the SP1 and the consequent absence of SP1-HDAC1 complex was already attributed behind the observed de-repression thereby triggering *GM2-synthase* transcription (18), the mechanism underlying the activation of the *GM2-synthase* gene was investigated. Among a number of candidate HATs (Histone acetyl transferases) identified in the two subsequent proteome profiling steps of enChIP-MS and CLASP-WB, p300 was previously reported to be a candidate enzyme for modulating both SP1 and SMAD 2, which is critical for regulating the stability and nuclear translocation (39) of the two proteins. Additionally, downstream analysis of data from the first enChIP-MS step not only identified p300 as one of the candidate HAT’s but further revealed a plausible association of p300 in complexes containing both SP1 and SMAD 2/4 (Fig. 3). In addition to this, the second CLASP-WB step also identified HAT-p300, SMAD2, 4 and SP1 to co-precipitate with *GM2-synthase* locus in three independent repeats (Fig. 4D & F) thereby further confirming the enChIP analysis and indicating a plausible role of p300 in *GM2-synthase* transcriptional regulation. To confirm this, we first explored the possibility whether at all any HATs might be involved in regulating *GM2-synthase* transcription. For this, low *GM2-synthase* expressing renal cancer cells, SK-RC-45 was treated with sodium butyrate (Nabu) either in the presence or absence of epigallactocatechin-3-gallate (EGCG), a pan-HAT inhibitor. EGCG was able to substantially diminish the effect of NaBu in *GM2-synthase* transcription (Fig. 8A), which indicated that under global HDAC inhibition, one of the HAT family members might be involved in the regulation of *GM2-synthase* transcription. To investigate the role of HAT-p300, p300 was knocked out from SK-RC-45 cells using CRISPR-Cas9, following which *GM2-synthase* activation was assessed upon NaBu treatment in both wild type SK-RC-45 cells and SK-RC-45 p300^-/-^cells. CRISPR-Cas9 mediated knockout of p300 resulted in a 5bp deletion at the exon 1 of p300 gene (Fig. 8B, lower panel) which caused a complete loss of p300 expression (Fig. 8B, upper panel). p300^-/-^knockout resulted in significant inhibition of NaBu mediated upregulation of *GM2-synthase* message levels, indicating a potential role of p300 in *GM2-synthase* transcription (Fig. 8C). This data was further reflected in HEK293T cells, where shRNA mediated knockdown of p300 resulted in significant abrogation of NaBu-mediated induction of *GM2-synthase* expression (Fig. 8D). Fig. 8E reflects the efficiency of shRNA-mediated knockdown of p300 expression in HEK293 cells by western blot, which shows a significant reduction in the p300 mRNA levels with respect to the non-transfected cells. Finally, over-expression of p300 in HEK-293T cells resulted in significant upregulation of NaBu-mediated *GM2-synthase* mRNA levels, as shown in Fig. 8F confirming the definite role of p300 in *GM2-synthase* transcriptional activation. Fig. 8G depicts an increase in the expression levels of *p300* mRNA which supports a positive correlation of p300 expression with *GM2-synthase* transcription upon NaBu treatment (55). Further Fig. 8H shows a time-dependent increase in p300 protein level in response to NaBu treatment as well as its activatory acetylation marks in line with previous reports (56). Surprisingly, Nabu treatment does not change the association of Ac-p300 to *GM2-synthase* promoter (Fig. 8I), suggesting that the activatory role of p300 in regulating *GM2-synthase* transcriptional activity is not exerted through changing the association levels of its active (acetylated) form with *GM2-synthase* promoter, and involves some other mechanisms possibly through its co-regulators.

**Fig 8.**
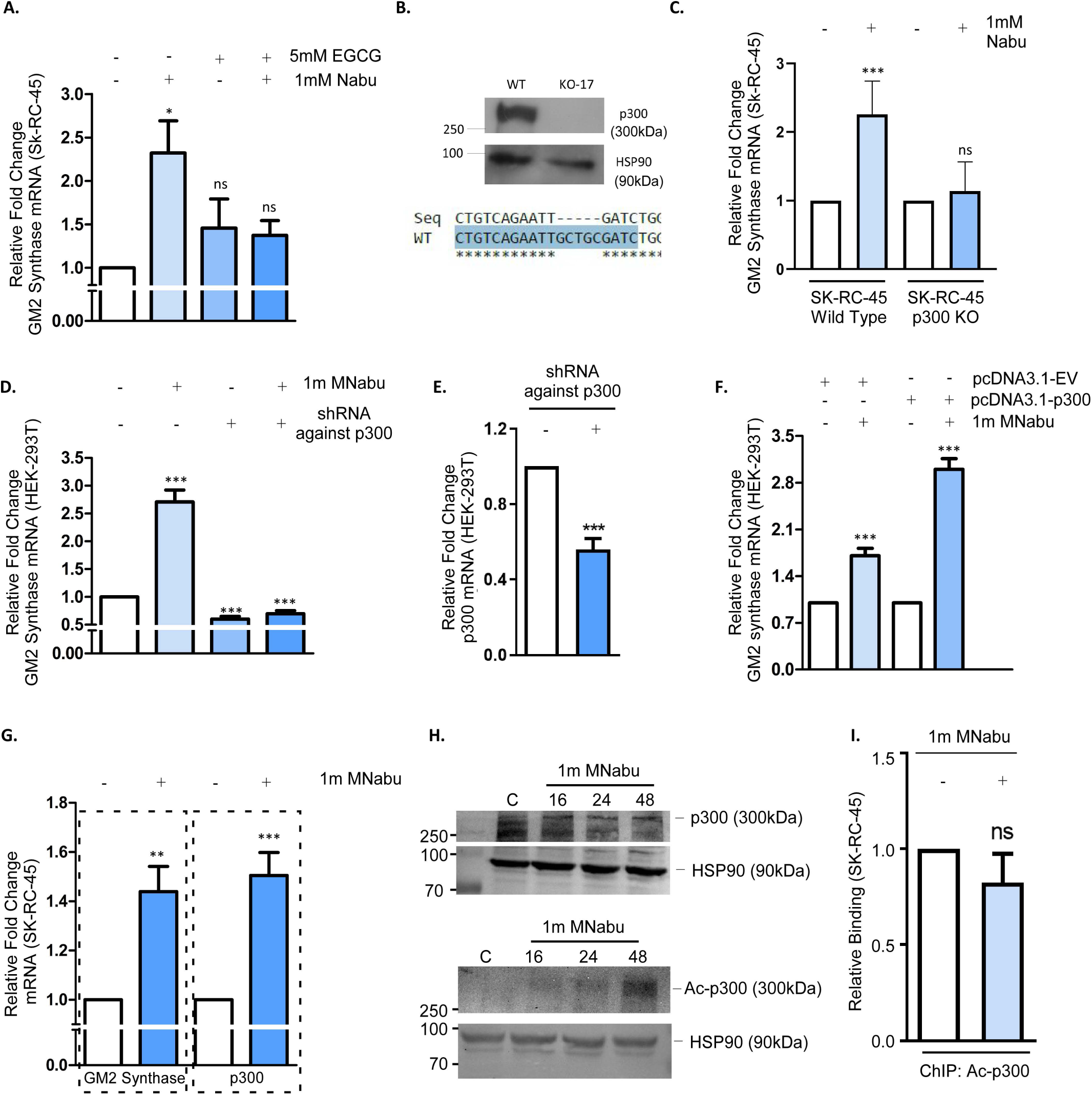
HAT-p300 acts as a transcriptional activator of GM2 synthase gene. A. Relative fold change of GM2 synthase mRNA upon global HDAC inhibition along with simultaneous global HAT inhibition, showing the involvement of HATs in GM2 synthase transcriptional regulation. B. Western blot validation and genomic characterization of p300 knock out in SK-RC-45 cells. C. Relative fold change of GM2 synthase mRNA in WT and p300 knockout SK-RC-45 cells upon HDAC inhibition. Statistical significance calculated with respect to the respective controls for both WT and KO17 cell lines. D&E. Relative fold change of GM2 synthase mRNA upon shRNA mediated knockdown of p300 in HEK-293T cells, and validation of the knockdown via RT PCR. F. Relative fold change of GM2 synthase mRNA upon p300 over-expression in HEK-293T cells. G. Relative fold change of GM2 synthase mRNA and p300 upon global HDAC inhibition. H. Western blots depicting the change of p300 protein levels (Top Panel) and the change in p300-acetylation levels (Bottom Panel) under global HDAC inhibition. I. Relative binding of Ac-p300 levels with GM2-synthase TSS in response to global HDAC inhibition. Error bars represent mean ±S.E.M of at least three independent determinations. (Student’s t test; *p 0.05**, p 0.01; ***, p 0.001). ns, not significant. All experiments are repeated with at least three biological replicates.

### HAT p300 regulates *GM2-synthase* transcription through regulation of SP1 and SMAD 2/4 Complex

The identification of p300 through the two-step proteome profiling pipeline and subsequent data establishing p300 as a positive regulator of *GM2-synthase* transcription prompted us to further investigate the possible role of p300 in the regulation of SP1 or SMAD 2/4 activity guiding *GM2-synthase* transcription. This hypothesis stemmed from the fact that p300 was already a well-known interacting member of both SP1 and SMAD 2 and known to modify both SP1 as well as SMAD 2 (38, 39). To find out whether at all, p300 has any role in NaBu-dependent degradation of SP1, time-dependent degradation of SP1 was assessed in both wild type as well p300 KO SK-RC-45 cells. While the wild type SK-RC-45 cells showed a time dependent decrease in SP1 protein levels as expected (18), the rate of decrease in SP1 expression levels was significantly slower in the p300 ^-/-^ SK-RC-45 cells (Fig. 9A & 9B), indicating that indeed p300 plays a definitive role in regulating the stability of the SP1 protein. To confirm whether the acetylation and ubiquitination status of SP1 proteins in response to NaBu treatment are also p300-dependent, both the wild type as well as the p300 KO cells were treated with NaBu, and SP1 was immunoprecipitated followed by western blot for looking at the acetylation and ubiquitination status of SP1. Data from Fig. 9C shows that while NaBu treatment caused increased SP1 acetylation as well as ubiquitination in the wild type SK-RC-45 cells, no significant acetylation or ubiquitination of SP1 was evident in the SK-RC-45 p300^-/-^ cells, indicating that indeed p300 was responsible for acetylation, ubiquitination resulting in subsequent lowering of cellular SP1 levels thereby activating *GM2-synthase* transcription. This was further reflected in a change of NaBu-dependent SP1 binding to the *GM2-synthase* promoter in wild type versus p300 KO cells (Fig. 9D). Data from Fig. 9D clearly shows that while NaBu causes a loss in SP1 association to the SP1 binding sites at +37 location of *GM2-synthase* promoter (ChIP) in wild type SK-RC-45 cells, no such loss of SP1 binding was apparent in the SK-RC-45 p300^-/-^ cells, clearly confirming that SP1 binding to the *GM2-synthase* promoter is p300-dependent. Next, the plausible role of p300 was explored in regulating activity of SMAD 2/4 which was found to activate *GM2-synthase* transcription (Fig. 5 and 6). NaBu mediated phosphorylation (Fig. 9E) of SMAD 2 was visibly reduced in the SK-RC-45 p300^-/-^ cells, indicating p300-dependent activation of SMAD 2. Time-dependent nuclear localization (Fig. 9F) of SMAD 2 & 4 in response to NaBu treatment in the p300 WT cells was significantly inhibited in the SK-RC-45 p300^-/-^ cells indicating that subsequent formation of the nuclear SMAD 2/4 nuclear complex under HDAC inhibition is critically regulated by p300. This indicates that p300 regulates *GM2-synthase* transcription upon global HDAC inhibition by acetylation and ubiquitination of SP1 and phosphorylating SMAD 2. While p300-dependent acetylation and ubiquitination of the repressor SP1 resulted in its degradation, phosphorylation of SMAD 2/4 complex led to its activation and nuclear translocation, ultimately resulting in transcriptional activation of *GM2-synthase* gene.

**Fig 9.**
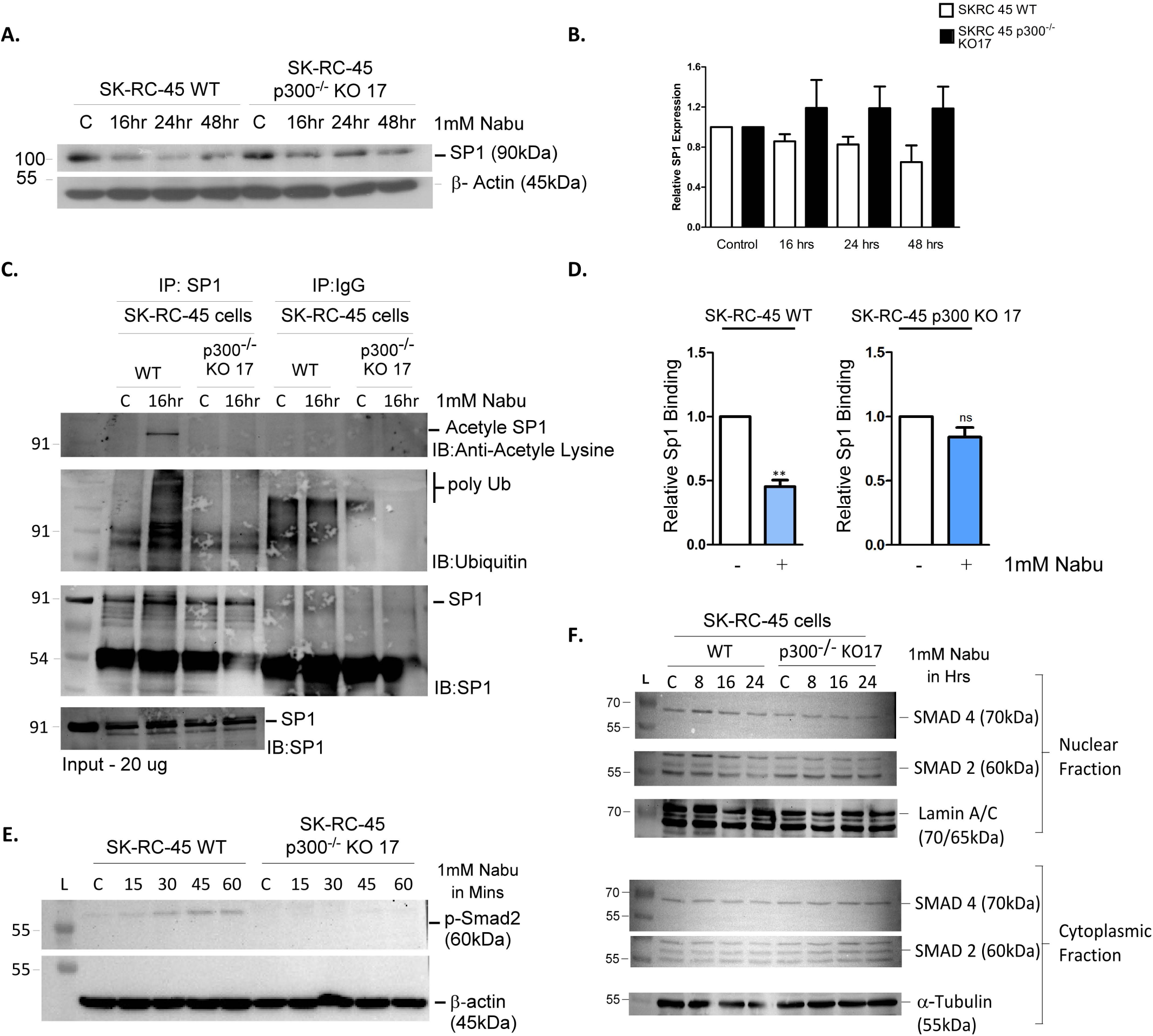
p300 positively regulates GM2 synthase promoter activity through its modulations on SP1 and SMAD proteins. A&B. Western blot showing the comparison between the time correlated decrease in SP-1 protein levels in both wild type and p300 knockout SK-RC-45 cells upon global HDAC inhibition. Graph at fig 9B only represents the quantification of the SP1 levels obtained in Fig 9A. C. Comparison between the acetylation and ubiquitination status of SP-1 in both wild type and p300 knockout SK-RC-45 cells upon global HDAC inhibition D. Relative binding of SP1 with its predicted binding sites upon global HDAC inhibition in both wild type and p300 knockout SK-RC-45 cells. E. Comparison between change in SMAD2 phosphorylation status in both wild type and p300 knockout SK-RC-45 cells upon global HDAC inhibition. F. change in SMAD2/4 nuclear localization status in both wild type and p300 knockout SK-RC-45 cells upon global HDAC inhibition. Error bars represent mean ± S.E of at least three independent determinations. (Student’s t test; *p 0.05**, p 0.01; ***, p 0.001). ns, not significant. Fig 9E was performed twice with all other experiments of this figure performed at least three times.

## Discussion

*GM2-synthase* is a critical regulator of ganglioside biosynthetic pathway and its aberrant over-expression in various cancers is well established (57, 58). In fact, GM2 over-expression has been linked with increased migration and invasion, epithelial to mesenchymal transition (EMT) and suppression of host immune response in cancer (5), while over-expression of *GM2-synthase*, at least in the case of renal cell carcinoma (RCC) has been associated with poor prognosis (14). However, despite its aberrant over-expression in several cancers, the mechanistic basis underlying the regulation of *GM2-synthase* expression has never been explored. Report from our laboratory first established an epigenetic mechanism underlying the transcriptional de-regulation of the *GM2-synthase* gene in RCC, where acetylation-dependent degradation of the transcriptional repressor, SP1 leading to reduced binding of the SP1-HDAC1 repressor complex on the *GM2-synthase* TSS was identified as a central mechanism for the observed over-expression of *GM2-synthase* in RCC. Although this report addressed some key questions including but not limited to the fact that withdrawal of the *GM2-synthase* repression upon acetylation-dependent proteasomal degradation of the SP1-HDAC1 complex leads to the observed over-expression of gene expression, the mechanism that poises the *GM2-synthase* transcriptional machinery is still not clearly understood. Hence, identification of a plausible “activator” complex holds paramount importance in obtaining clarity in the underlying transcriptional complexity of *GM2-synthase* expression. Although Co-IP followed by mass spectrometric analysis of the immunoprecipitated proteome was utilized traditionally for identification of the proteome, however these approaches lacked the specificity required to understand the association of the players at the genomic context. With the advent of CRISPR-Cas technology and utilizing the DNA-binding ability of a modified Cas9 protein, namely dCas9 (dead Cas9) lacking the DNA-cleaving activity, the proteome in the context of the specific genomic locus of interest is identifiable using enChIP (engineered DNA binding molecule mediated chromatin immunoprecipitation) and CLASP (Cas9 locus-associated proteome) techniques (19, 59, 60). With the primary objective of pulling down the proteome associated with the TSS of the *GM2-synthase* gene, this study initiates with profiling the entire proteome associated with the *GM2-synthase* promoter using enChIP-MS and CLASP-WB as two steps, followed by characterisation and identification of the possible activators of *GM2-synthase* transcription. This study further ventures towards establishment of SMAD2/4 as an activator and how activation by SMAD 2/4 play hand-in-hand with the repression mediated by SP1 in modulating *GM2-synthase* transcription. The study additionally establishes p300 as a central player positively regulating *GM2-synthase* transcription through its ability to phosphorylate and activate SMAD 2/4 while degrading SP1 proteins thereby regulating their respective activities.

For profiling the promoter associated proteome we designed a two-step locus specific proteome profiling regime by taking help of two previously reported CRISPR Cas9 techniques, both of which targets a catalytically inactive dCas9 to the locus of interest using a guide RNA followed by its immunoprecipitation to co-purify the proteins present at the locus. Following the successful targeting as evidenced from the enrichment of the targeted locus (Fig. 1F), we performed mass spectrometric analysis to identify the components of *GM2-synthase* promoter associated proteome as the first hypothesis generating step to nominate pool of regulators to be further validated by the second CLASP-WB step (Fig. 4), before challenging the obtained findings through subsequent layers of orthogonal validation. The components of the promoter associated proteome identified through the mass spectrometric analysis revealed a diverse group of proteins ranging from histone modifiers to chromatin architectural elements. While the presence of histone modifiers and transcription factors were of particular interest because of their direct relation with regulation of gene expression, the other identified components such as CTCF or BRD 4 opened up the possibilities of long-range DNA interactions or enhancers (61–63) to also have an impact on *GM2-synthase* transcription. Also, within the group of histone modifiers, presence of proteins such as lysine demethylase and methyl transferases opened up the possibilities of histone methylation marks to control *GM2-synthase* transcription, none of which has been observed before. While our previous publication (18) revealed that there is no correlation between *GM2-synthase* transcriptional regulations with variations with CPG island methylation patterns of its promoter, the identification of various regulators which utilize the CPG islands as probable docking sites, in the proteome profiling demanded further investigation about the possibility of an indirect regulatory impact (64), although this is beyond the scope of the present study.

In the second step of the two-step proteomic profiling regime, we systematically isolated the *GM2-synthase* locus using, purified dCas9 and *in-vitro* synthesised guideRNA and detected various important proteins nominated through the initial enChIP-MS step of this two-step regime. In total 11 important regulators, shortlisted from the first enChIP step namely, SP1, SMAD2, SMAD4, HDAC1, STAT3, p53, p65, p300, c-JUN, TEAD-1 and SMARCE1 were repeatedly detected to co-purify with *GM2-synthase* locus, while IKBα remained undetected acting as a negative control.

The two-step dCas9 mediated proteomic profiling pipeline identified various potential activators of transcription among which SMAD 2/4 had a predicted binding site at *GM2-synthase* promoter. Since, SMAD 2/4 is already recognized as a mediator of TGF β signalling, we wanted to see whether TGF β could at all influence *GM2-synthase* expression (Fig. 5A), which was confirmed by a significant upregulation of *GM2-synthase* message level, along with an even more amplified up-regulation of ganglioside GM2 levels (Fig. 5B & 5C). To further investigate the possible involvement of SMAD 2/4 in *GM2-synthase* transcriptional regulation we subjected low *GM2-synthase* expressing cell lines with SMAD 2/4 over-expression (Fig. 5D). Under SMAD over-expression as well, our studies found a strong co-relation of SMAD 2/4 activity and *GM2-synthase* transcription as well as an amplified up-regulation in ganglioside GM2 levels (Fig. 5E&S3D). Additionally, luciferase experiments with either deletion and/or mutation constructs revealed that the proximal SMAD binding element and SP1 binding sites had contrasting roles. While the SP1 binding site at -311/-295 like the previously reported site at +109/+125 also acted as a repressor of transcription, the proximally present SMAD binding element at -324/-318 was found to act as an activator and that the SP1 binding site interfered with the activity of the SMAD binding element in a way that the presence of the SP1 binding site resisted the inducibility of the SMAD binding element. In connection to this finding, *in-silico* modelling studies done with the Smad-SP1-DNA complex (Fig. 7A and B) also revealed that the simultaneous binding of SMAD and SP1 with their immediate adjacent DNA binding sites to be energetically un-favourable due to steric hindrance. Therefore, this modelling together with preliminary check of their stabilities rationalized the experimental message coming out from the luciferase promoter experiments (Fig. 6C, D and E), that the presence of SP1 binding site at - 311/-295 hinders the inducibility of SMAD binding element positioned at just 6 nucleotides downstream at -324/-318 in the context of *GM2-synthase* transcription.

The formation of the SMAD 2/4 complex at the *GM2-synthase* TSS was indicated by increased binding of SMAD 2/4 at the TSS (Fig. 5G) in response to NaBu treatment, indicating a plausible involvement of SMAD 2/4 in the transcriptional regulation of the *GM2-synthase* gene. Additionally, changes in phosphorylation status of SMAD 2 in response to NaBu treatment confirms SMAD 2 activation, which is a pre-requisite for its nuclear translocation, which is yet again confirmed by increased localization of SMAD 2/3 in the nucleus time-dependently (Fig. 5J-K).

As our previous study reported the involvement of a HDAC1 in regulating the transcriptional activity, we wanted to investigate the involvement of HATs in *GM2-synthase* transcription. The presence of various HATs including p300 in the two-steps of the proteomic profiling pipeline also prompted us to test our hypothesis further. To investigate the involvement of HATs we subjected low *GM2-synthase* expressing renal cancer cell lines to global HAT inhibition on a HDAC inhibited background. The experimental demonstration that HDAC inhibition did not lead to the up-regulation of *GM2-synthase* transcription under global HAT inhibition, provided us the evidence that one of the HAT family members were responsible for *GM2-synthase* transcriptional up-regulation when HDACs were inhibited. As both SMAD2 and SP1 were previously reported to be interacting partners of p300, and as p300 is already reported to control SMAD and SP1s activity (65, 66), we knocked out p300 from SK-RC-45 cells and subjected the cells to global HDAC inhibition. This also resulted in abrogation of HDAC inhibitor mediated up-regulation of *GM2-synthase* transcription confirming the hypothesis of p300’s involvement in *GM2-synthase* transcription. This finding was further supported by our shRNA mediated knockdown and p300 over-expression experiments which revealed p300 to be a positive regulator of *GM2-synthase* transcription. In addition to the previous findings, p300 itself showed a transcriptional up-regulation upon global HDAC inhibition, which co-related with the *GM2-synthase* transcriptional up-regulation (Fig. 8G). Although p300s transcriptional up-regulation by global HDAC inhibition was reported earlier as well in prostate cancer cell lines (55), the correlation between the HDAC inhibition mediated transcriptional up-regulation of both p300 and *GM2-synthase* added another important angle to our finding that under situations when HDACs are less active compared to HATs, not only the regulatory void is filled up by p300 molecules already present in the system but also causes an increase of p300 levels itself both in terms of mRNA and protein levels, as well as p300 acetylation status, leading to an even amplified effect of p300 on *GM2-synthase* transcriptional regulation. But the mechanism underlying such an upregulation of p300 expression in the context of *GM2-synthase* transcription needs detailed exploration which falls outside the scope of this study. Astonishingly, although HDAC inhibition caused an induction of p300 mRNA and protein levels in addition to an increase in its acetylation levels, the binding of Ac-p300 to the *GM2-synthase* locus, in response to global HDAC inhibition did not change (Fig. 8I), indicating that the positive effects exerted by p300 on *GM2-synthase* gene transcription, does not take place by changing the association status of its active form with *GM2-synthase* TSS.

To find out the inter-relation of SMAD-SP1 with p300 in the context of *GM2-synthase* transcription, we investigated the impact of p300s absence on the regulatory behaviours of SMAD and SP1 uncovered by our previous experiments. Surprisingly, the degradation of repressor SP1 under the influence of global HDAC inhibition, which provided the primary mechanistic insight to *GM2-synthase* transcriptional regulation (18), was completely absent in p300 knock out SK-RC-45 cells, confirming a pivotal role of p300 in maintenance of the stability of SP1 inside the cells. This finding was further reflected in the binding status of SP1 with the +37/+187 region of the promoter of interest, while the wild type cells showed a decrease in SP1 binding to the locus in response to global HDAC inhibition, this change in SP1 binding was absent in p300 knock out cells. These data strongly imply that SP1 may be a strong candidate for acetylation and ubiquitination by p300, since SP1 acetylation as well as subsequent ubiquitin-dependent degradation has been previously reported under global HDAC inhibition. This was confirmed by the acetylation status of SP1 under global HDAC inhibition which shows that HDAC inhibition mediated acetylation of SP1 was also happening through p300. Collectively these data present an important aspect of transcriptional regulation of *GM2-synthase* gene and the role of SMAD-p300-SP1 axis therein. This study highlights the HAT, p300 as critical player in reciprocal regulation of the activities of SMAD 2 and SP1 through phosphorylation and acetylation respectively, which activates SMAD and targets SP1 for degradation, thereby poising *GM2-synthase* for transcriptional activation. In totality, this study started with a two-step dCas9 based proteomic profiling regime, where the initial enChIP-MS and its downstream analysis step played the role of hypothesis generation, nominating important proteins to the second dCas9 mediated CLASP-WB step, information obtained from which were deciphered layer by layer with various levels of orthogonal validations to ultimately decipher the SMAD-p300-SP1 axis in transcriptional regulation of *GM2-synthase* gene. Data from this manuscript demonstrate that GM2 synthase is regulated by a buffered SMAD dependent mechanism which is extremely sensitive to even slightest of perturbations at the transcriptional level and results in a much-amplified biochemical output, in terms of ganglioside GM2 levels. We further establish that this buffered inducibility results from the presence of a proximal SP1 binding element and deletion of this element restores the expected magnitude of transcriptional induction. Together these findings show, that it is the locus specific regulatory architecture rather than the absence of SMAD mediated regulation that explains this discrepancy between limited transcriptional induction and enhanced biochemical output. We also show that expression and activation of HAT-p300 correlates with GM2 synthase transcription, and it reciprocally regulates the degradation of repressor SP1 and activator SMAD, by regulating the acetylation mediated degradation of the former while inducing the activation and nuclear translocation of the later while this function of p300 is independent of the binding status of its active form to the GM2 synthase locus.

Further, this study implicates critical understanding of the molecular events occurring at the *GM2-synthase* promoter, as well as its significance in cancer. Along with p300 various other DNA-looping proteins and enhancer binding proteins came up in our mass spectrometric analysis such as BRD 4, CBP etc, and hence the possibility of distal DNA interactions as well as three dimensional events occurring at the locus remains an area to be ventured. Also, the dynamics of opening and closing of the chromatin with respect to the change in expression of *GM2-synthase*, remains to be explored. Although interesting, both these studies are beyond the scope of the current one and demand further investigation.

## Author contributions

K.B and S.B conceptualised the study and designed the experiments. S.B performed the experiments. A.B generated the p300 knock out cells, and provided critical as well as intellectual inputs from time-to-time. S.R performed immunostainings presented at Fig. 5B and Fig. 5E. B.W performed the mass spectrometry and analysed the mass spectrometry data. S.B, A.R, S.R and K.B analysed the data. S.G.D. and D.P. performed the *in-silico* modelling experiments. S.B and K.B wrote and edited the manuscript. K.B. conceived the project and provided overall supervision of the study.

## Acknowledgements

This work was supported by funds from the Indian Council of Medical Research (ICMR), New Delhi, Sanction number: 2019-0137-CMB/adhoc/BMS; Department of Science and Technology (DST), Sanction number: EMR/2016/001983, and CRG/2021/004623. Mr. Sounak Banerjee acknowledges University Grants Commission (UGC) for his fellowship.

## Data availability

The proteomics data generated in this paper is available with accession number PXD065551 in the Proteome Exchange Consortium via PRIDE partner repository.

---

**Graphical abstract. Proposed model on the role of SP1-HDAC1-p300-SMAD 2/4 regulatory axis on GM2 synthase transcription.** P300 regulates both SP1 and SMAD 2/4 complex to de-repress GM2 synthase transcription in high GM2 synthase expression conditions

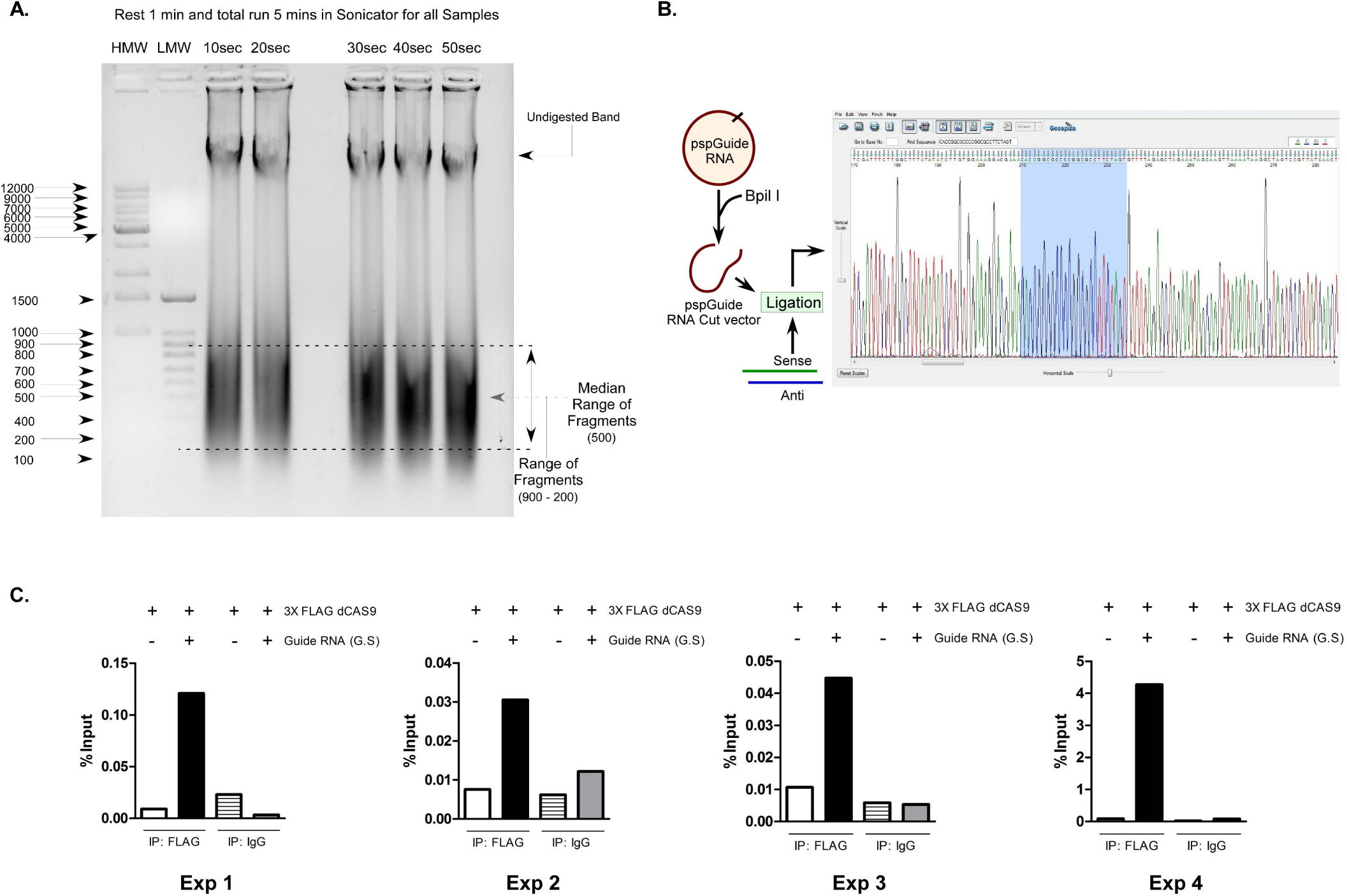

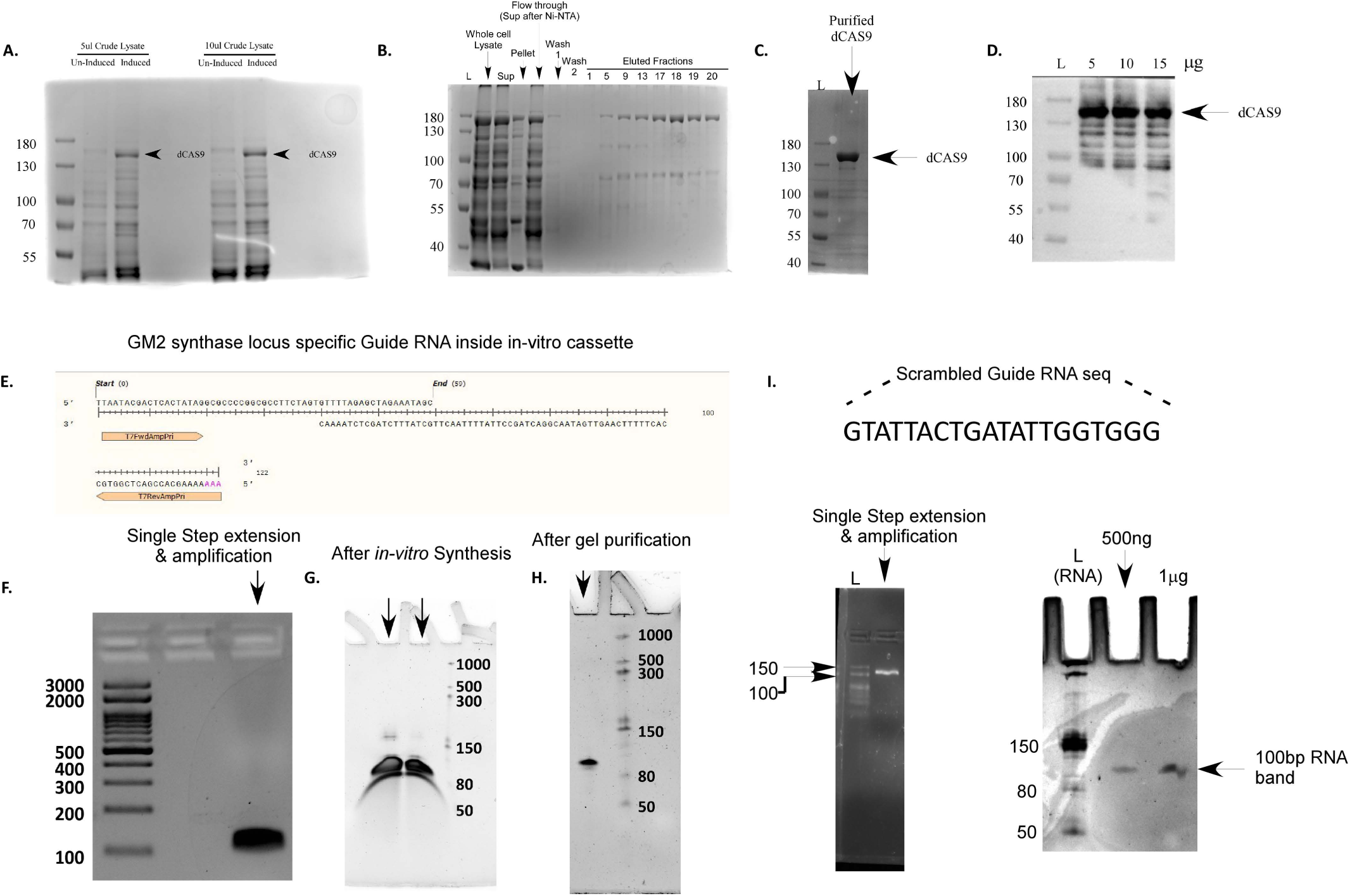

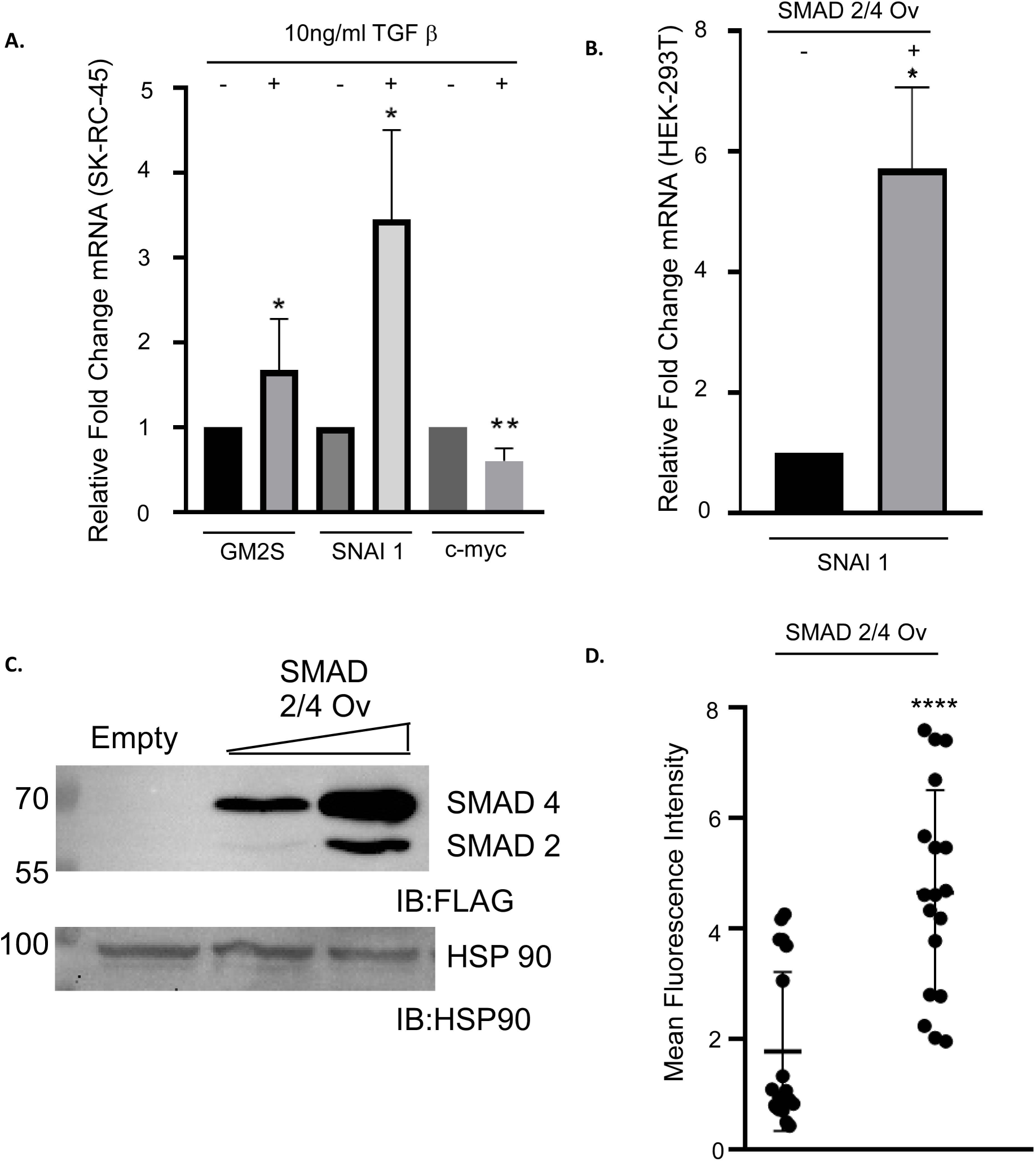

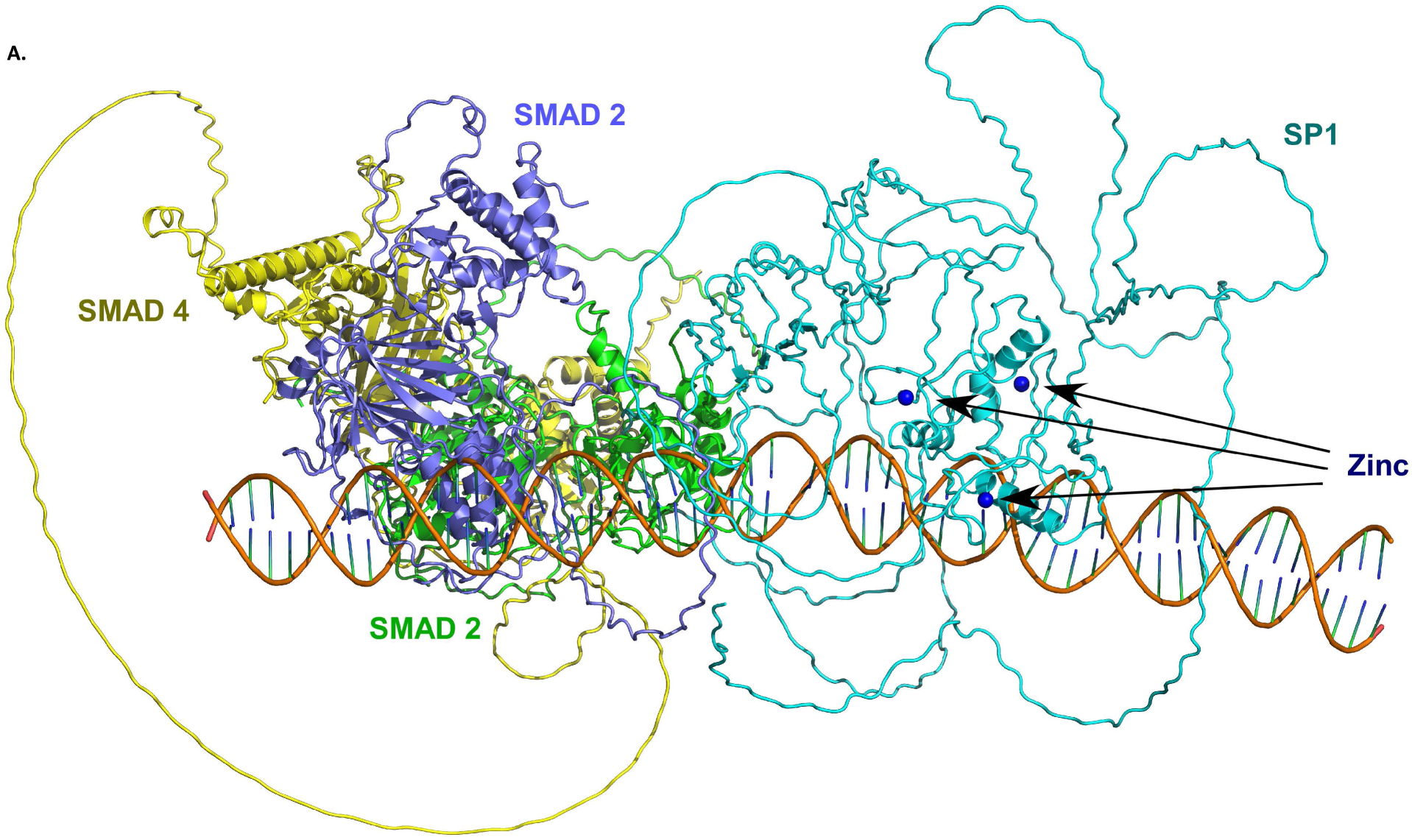

