## Supplementary Information Text for "dCas9 targeted proteome profiling reveals p300-mediated reciprocal regulation of SMAD and SP1 as a driver of GM2synthase transcription in renal cell carcinoma"

### dCas9 mediated targeted proteome profiling reveals SMAD-p300 axis as the transcriptional activator of GM2 synthase gene in Renal cell carcinoma.

Sounak Banerjee[1], Avisek Banerjee[1, 2#], Subha Ray[1#], Aishwarya Ray[1], Debarati Paul[1], Shubhra Ghosh Dastidar[1], Belinda Willard[3] and Kaushik Biswas*[1]

[1]Department of Biological Sciences, Bose Institute, Kolkata, West Bengal, 700091, India, [2]Department of Zoology, Ramakrishna Mission Vidyamandira, Belur Math, Howrah, 711202, .India, [3]Proteomics and Metabolomics SLR, Lerner Research Institute, Cleveland Clinic, Cleveland, OH, USA, 44195.

* To whom all correspondence should be addressed:

[#] These authors have contributed equally to this work.

Kaushik Biswas, Department of Biological Sciences, Bose Institute, Kolkata, EN-80, Bidhan Nagar, Salt Lake, Sector-V, Kolkata, West Bengal, 700091, India,

**Running Title**

Mechanistic dissection of GM2 synthase transcriptional regulation reveals SMAD-p300-SP1 axis

**List of supporting material**

1. Table S1 – List of oligo-nucleotides and primers.
2. Table S2 - Matrix scores of TFind webtool prediction of the SMAD binding element.
3. Supporting information Fig. S1.
4. Supporting information Fig. S2.
5. Supporting information Fig. S3.

#### 1. Supporting Information Table S1 – List of oligo-nucleotides and primers

**For Real-Time PCR**

| | **Forward** | **Reverse** |
|---|---|---|
| GM2-synthase | 5' –TTT GAC CCT GCA GAG CTG-3' | 5' –CTG AAC TTC CAC ACC CTG TAG-3' |
| GAPDH | 5' –ACA ACT TTG GTA TCG TGG AAG G– 3' | 5' –GCC ATC ACG CCA CAG TTT C– 3' |
| p300 | 5' – AGCCAAGCGGCCTAAACT C– 3' | 5' – TCACCACCATTGGTTAGTCC C– 3' |

**For enChIP, CLASP and ChIp experiments**

| **For ChIP Realtime PCR** | **Forward** | **Reverse** |
|---|---|---|
| GM2 Synthase -113 to +37 | 5'- AGCAGGCAATGGAATGGA TG-3' | 5'- AGCGCCCCGGCGCCTTCTA G-3' |
| GM2 Synthase +38 to +187 | 5'- CGCATTCCCCGCGCGGAGC - 3' | 5'- AGCCCCGGGGCAAAGCCG GG-3' |
| **For ChIP Semiquantitative PCR** | | |
| GM2 Synthase -363 to -203 | 5'- GAAATAGGGCCTTCAATTA C -3' | 5'- TTAGGATGAAGATAAAGG AA-3' |

| | |
|---|---|
| Guide enChIP Sense | 5'- CACCGCCTACAAGCTCAGAACGAGC-3' |
| Guide enChIP anti-Sense | 5'- AAACGCTCGTTCTGAGCTTGTAGGC-3' |

**For in-vitro guide RNA synthesis used as a part of CLASP experiments**

| | **Forward** | **Reverse** |
|---|---|---|
| T7 Amp primer | 5'- TAATACGACTCACTATAG-3' | 5'- AAAAAAAGCACCGACTCGGTGC-3' |

| | |
|---|---|
| Guide Sense For CLASP | 5'- TTAATACGACTCACTATAGCCTACAAGCTCAGAACGAGCGTTTTAGAGCTAGAAATAGC-3' |
| Guide Anti for CLASP | 5'- AAAAGCACCGACTCGGTGCCACTTTTTCAAGTTGATAACGGACTAGCCTTATTTTAACTTGCTATTTCTAGCTCTAAAAC-3' |

**For p300 targeting shRNA experiments**

| | |
|---|---|
| EP300 shRNA - Sense | 5'- CCGGCCAGCCTCAAACTACAATAAACTCGAGTTTATTGTAGTTTGAGGCTGGTTTTTG-3' |
| EP300 shRNA - Anti | 5'- AATTCAAAAACCAGCCTCAAACTACAATAAACTCGAGTTTATTGTAGTTTGAGGCTGG-3' |

**For Site Directed Mutagenesis**

| | |
|---|---|
| For SMAD SDM Wt Background For | 5'- TGCCCCCCATTATTTAACTTTTACTCAAGAGAAATCTAGA-3' |

| | |
|---|---|
| For SMAD SDM Wt Background Rev | 5'-TCTAGATTTCTCTTGAGTAAAAGTTAAATAATGGGGGGCA-3' |
| For SMAD SDM on SP1 SDM Background For | 5'-TATTTCCCATTATTTAACTTTTACTCAAGAGAATCTAGAT-3' |
| For SMAD SDM on SP1 SDM Background Rev | 5'-ATCTAGATTCTCTTGAGTAAAAGTTAAATAATGGGAAATA-3' |

**For CRISPR mediated knockout of p300 in SK-RC-45 cells**

| | |
|---|---|
| EP300_Sense | 5'-CACCGCTGTCAGAATTGCTGCGATC-3' |
| EP300_Antisense | 5'-AAACGATCGCAGCAATTCTGACAGC-3' |

2. **Supporting Information Table S2 - Matrix scores and p-value of TFind webtool prediction of the SMAD binding element.**

| Position | Sequence | Rel Score | Score | Rel Score Adj | Score Adj | p-value | Strand |
|---|---|---|---|---|---|---|---|
| 840 | gagTGTCTGtta | 1.000000 | 7.931569 | 1.000000 | 8.313832 | 2.590000e-04 | - |

3. **Supporting Information Table S3 - Matrix scores and p-value of TFind webtool prediction of the SP1 binding element.**

| Position | Sequence | Rel Score | Score | Rel Score Adj | Score Adj | p-value | Strand |
|---|---|---|---|---|---|---|---|
| 863 | gcaGGGCGGgga | 1.000000 | 7.931569 | 1.000000 | 6.945518 | 5.090000e-04 | + |

#### 4. Supporting information S1.

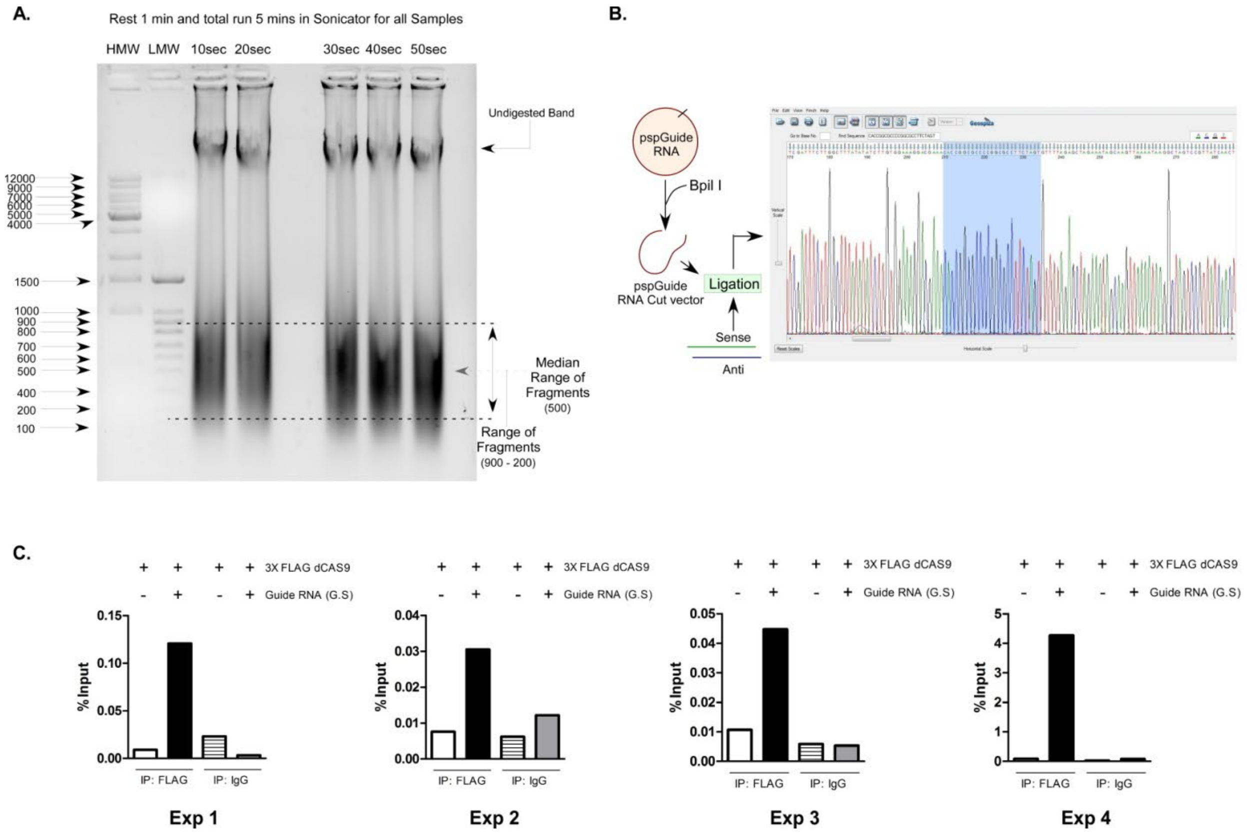

*Fig S1 - A. Fragmentation of isolated chromatin for enChIP – MS and size selection of chromatin fragment with 500 bp median length. B. cloning of gRNA targeting GM2 synthase promoter (-131/-111) into guideRNA expression vector (Bbs1 ver2, Addgene-85586). C. Standardization of specific isolation of GM2 synthase locus with enChIP technique.*

#### 5. Supporting information S2.

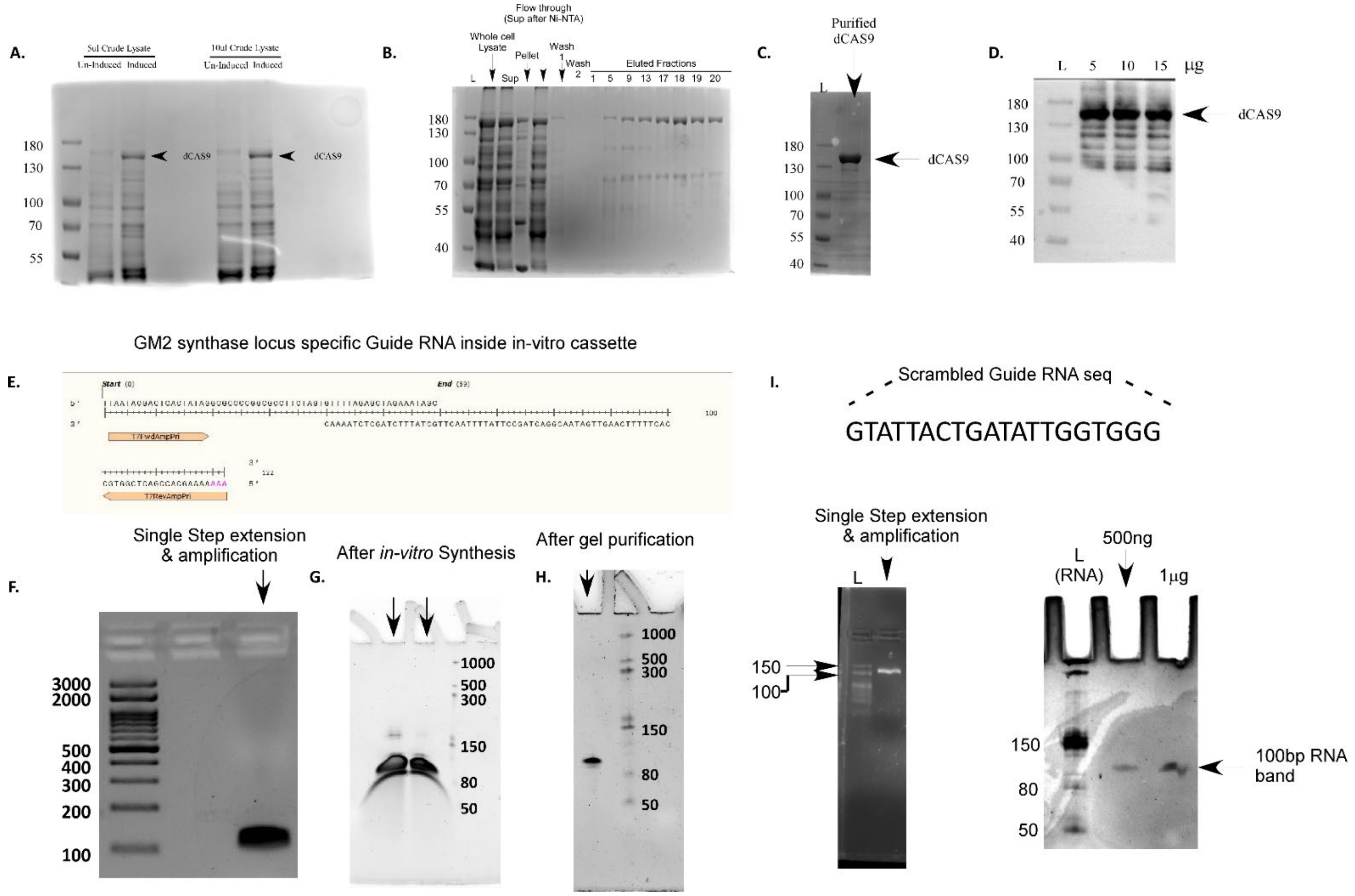

*Fig S2 - A. Expression of dCas9 from pCT310-dCas9 expression vector in e.coli upon IPTG induction. B. Ni-NTA purification of dCas9 from e.coli transformed with pCT310. C&D. Coomassie gel and western blot (anti-FLAG) of purified dCas9 after size exclusion chromatography. E. Oligo-nucleotide sequences annealed for single step overhang extension PCR for making dsDNA template for in-vitro gRNA synthesis. F. dsDNA template after overhang extension PCR. G. Total product of in-vitro RNA synsthesis reaction run on a 6% PAGE containing 7M urea. H. purified gRNA after gel elution from previous step (FigS2G), run on a 6% PAGE containing 7M urea. I. Synthesis of control gRNA for CLASP-WB step. Sequence of unique oligo for control gRNA synthesis (Top Panel), dsDNA template after single step overhang extension PCR (Bottom left panel) and purified gRNA after gel elution from previous step, run on a 6% PAGE containing 7M urea (Bottom right panel).*

**5. Supporting information S3.**

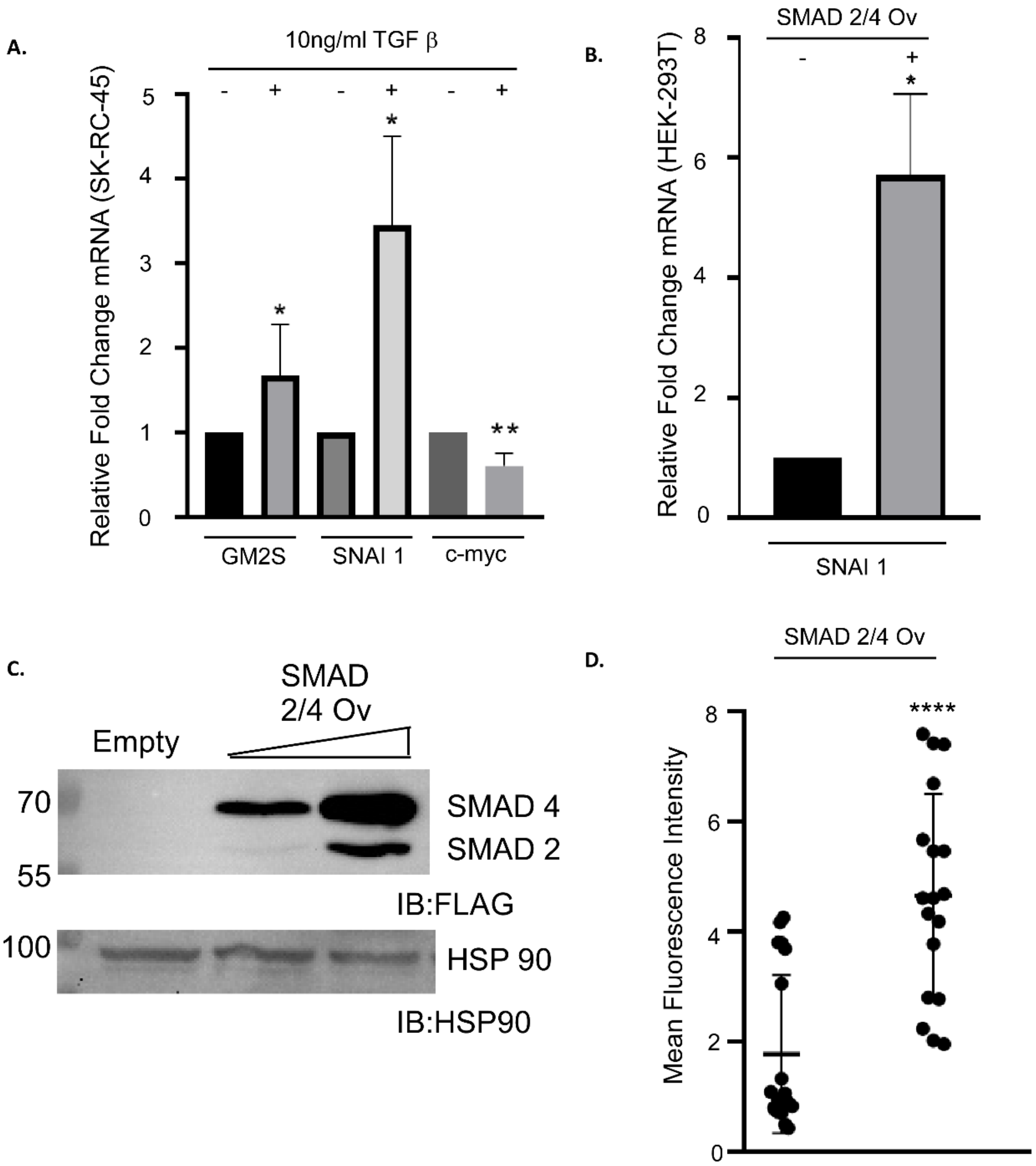

*Fig S3 - A. Changes in mRNA levels for SMAD target genes SNAI1 and c-myc in comparison to GM2-synthase gene upon TGF β treatment. B. Changes in mRNA levels for SMAD target gene SNAI1upon SMAD2/4 over-expression. C. Protein levels of SMAD2 and SMAD4 upon SMAD2/4 over-expression in HEK-293T cells. D. Quantification of mean fluorescence intensity in HEK-293T cells over-expressed with SMAD 2 and 4 compared to control cells.*

#### 6. Supporting information S4

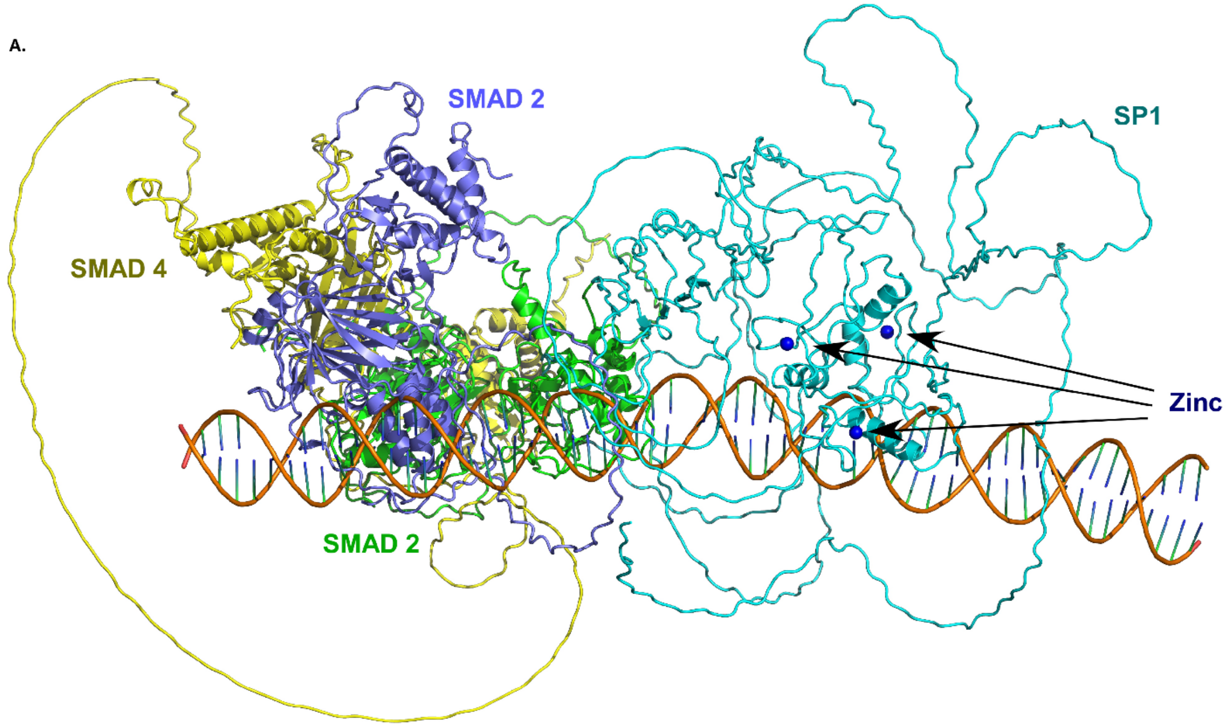

*Fig S4 - A. Cartoon representation of the full DNSPAD complex. SMAD2 monomers 1 and 2 are shown in green and lavender, respectively, while SMAD4 is colored yellow. The DNA double helix is depicted in orange, SP1 is represented in cyan and zinc ions are represented as blue spheres.*
